# Unprecedentedly efficient CUG initiation of an overlapping reading frame in *POLG* mRNA yields novel protein POLGARF

**DOI:** 10.1101/2020.03.06.980391

**Authors:** G Loughran, AV Zhdanov, MS Mikhaylova, FN Rozov, PN Datskevich, SI Kovalchuk, MV Serebryakova, S Kiniry, AM Michel, PBF O’Connor, DB Papkovsky, JF Atkins, PV Baranov, IN Shatsky, DE Andreev

## Abstract

While near cognate codons are frequently used for translation initiation in eukaryotes, their efficiencies are usually low (<10% compared to an AUG in optimal context). Here we describe a rare case of highly efficient near cognate initiation. A CUG triplet located in the 5’ leader of *POLG* mRNA initiates almost as efficiently (~60-70%) as an AUG in optimal context. This CUG directs translation of a conserved 260 triplet-long overlapping ORF, which we call *POLGARF* (*<u>POLG</u>* <u>A</u>lternative <u>R</u>eading <u>F</u>rame). Translation of a short upstream ORF 5’ of this CUG governs the ratio between DNA polymerase and POLGARF produced from a single *POLG* mRNA. Functional investigation of POLGARF points to extracellular signalling. While unprocessed POLGARF resides in the nucleoli together with its interacting partner C1QBP, serum stimulation results in rapid secretion of POLGARF C-terminal fragment. Phylogenetic analysis shows that *POLGARF* evolved ~160 million years ago due to an MIR transposition into the 5’ leader sequence of the mammalian *POLG* gene which became fixed in placental mammals. The discovery of *POLGARF* unveils a previously undescribed mechanism of *de novo* protein-coding gene evolution.

**Significance Statement:** In this study, we describe previously unknown mechanism of *de novo* protein-coding gene evolution. We show that the *POLG* gene, which encodes the catalytic subunit of mitochondrial DNA polymerase, is in fact a dual coding gene. Ribosome profiling, phylogenetic conservation, and reporter construct analyses all demonstrate that *POLG* mRNA possesses a conserved CUG codon which serves as a start of translation for an exceptionally long overlapping open reading frame (260 codons in human) present in all placental mammals. We called the protein encoded in this alternative reading frame POLGARF. We provide evidence that the evolution of *POLGARF* was incepted upon insertion of an MIR transposable element of the SINE family.

## Introduction

The process of translation can be described in four steps: initiation; elongation; termination and ribosome recycling. It is believed that protein synthesis is mostly regulated at the level of initiation. In eukaryotes, the scanning model for translation initiation postulates that the small ribosomal subunit, in complex with initiation factors and Met-tRNAi, enters at the 5’ end of mRNA and then scans towards the 3’ end (1). Base-pairing interactions between the anticodon of the Met-tRNAi and an AUG codon in the mRNA halts ribosome scanning and sets the reading frame for subsequent elongation steps (2). Notably, due to mRNA mispairing with the anticodon of Met-tRNA_i_, initiation can also occur at most triplets that differ from AUG by a single nucleotide (near-cognate), albeit with much lower efficiency (3).

Initiation efficiency on any translation initiation site (TIS) critically depends on its surrounding nucleotide context. In pioneering work, Kozak proposed that the context comprising 6 nt before and 1 nt immediately following a potential initiation codon has significant influence on the recognition of an initiation site (4). In agreement with Kozak, recent high-throughput analysis of all possible initiation contexts revealed RYMRMV<u>AUG</u>GC as the optimal context in human and mouse cells and additionally revealed synergistic effects of neighboring nucleotides (5).

TISs in unfavorable context can be bypassed by the scanning ribosome in a process known as leaky scanning. Since many mammalian mRNA 5’ leaders possess AUG codons, as well as many potential near-cognate start codons, then leaky scanning must be widespread. However, the mere presence of a potential TIS in a 5’ leader doesn’t necessarily guarantee initiation there. Until recently, it was difficult to estimate how frequently upstream TISs (uTISs) are recognized by scanning ribosomes in living cells. This can now be directly addressed since the emergence of the ribosome profiling technique (riboseq), which allows monitoring of global translation at single nucleotide resolution (6). Riboseq revealed widespread translation in the 5’ leaders of mRNAs, especially in mammalian cells (7–9).

What is the role of translation initiation in 5’ leaders? In some instances it gives rise to N-terminal extensions (10–12) though in most cases they result in translation of short (some are simply AUG-stop) upstream open reading frames (uORFs). While it is believed that most uORFs suppress translation of their main protein-coding sequence (13–15), a number of uORFs are involved in more specialized regulation of translation, ranging from selective stress responses to eIF2 phosphorylation (16–19), to metabolite sensing (20–22).

Our attention was drawn to one such uORF within a mRNA encoding a catalytic subunit of mammalian mitochondrial DNA polymerase (POLG). POLG is a hotspot for more than 200 known mutations in humans that cause mitochondria-associated diseases such as PEOA1, SANDO, AHS, and MNGIE (https://tools.niehs.nih.gov/polg/). Disease development is believed to result from a gradual depletion of mtDNA due to polymerase dysfunction(s). Transgenic mice with a mutated POLG causing proofreading deficiency (Polg mutator mouse) develop a mtDNA mutator phenotype, which is characterized by low mtDNA copy number, decreased lifespan and premature ageing (23).

There is a single AUG within the *POLG* 5’ leader which is expected to initiate translation of a conserved 23 codon uORF. Here we show that, contrary to expectations, the removal of the upstream AUG suppresses translation of the *POLG* coding sequence. Exploring the unusual effect of the uORF mutation revealed highly efficient CUG initiation of a 260-codon long alternative reading frame (−1) overlapping the *POLG* main ORF. Thus, the *POLG* mRNA turns out to be a dual coding messenger.

## Results

### A CUG codon located upstream of POLG CDS governs translation of a long overlapping reading frame that encodes POLGARF

The 5’ leader of the POLG mRNA contains a 23-codon conserved AUG-initiated upstream uORF. To determine whether translation of this uORF affects the synthesis of POLG we fused the whole 5’ leader of *POLG* (plus 33 nt downstream of the main ORF start codon) to a Firefly luciferase (Fluc) reporter and explored the effect of preventing uORF translation on reporter activity (Fig. 1A). In general, uORF translation represses translation of the main ORF by decreasing the number of scanning 43S preinitiation complexes that reach the main ORF start codon (15), however, here, translation of the uORF enhances main ORF translation (Fig. 1A). In search for potential explanations we examined publicly available ribosome profiling data (24). Within the *POLG* mRNA, the phase of triplet periodicity of ribosome footprints supports translation of an alternative reading frame (frame −1) that overlaps the *POLG* CDS (Fig. 1B). This footprint density is higher than the density of footprints aligning to the *POLG* CDS and decreases abruptly at the first stop codon in the −1 frame located in exon 3 of *POLG* (Fig. 1C). Notably, this −1 frame stop codon is universally conserved across placental mammals (Fig. 1C, middle panel) and the pattern of synonymous substitutions in the *POLG* reading frame before the −1 frame stop codon suggests dual coding in this region.

**Figure 1.**
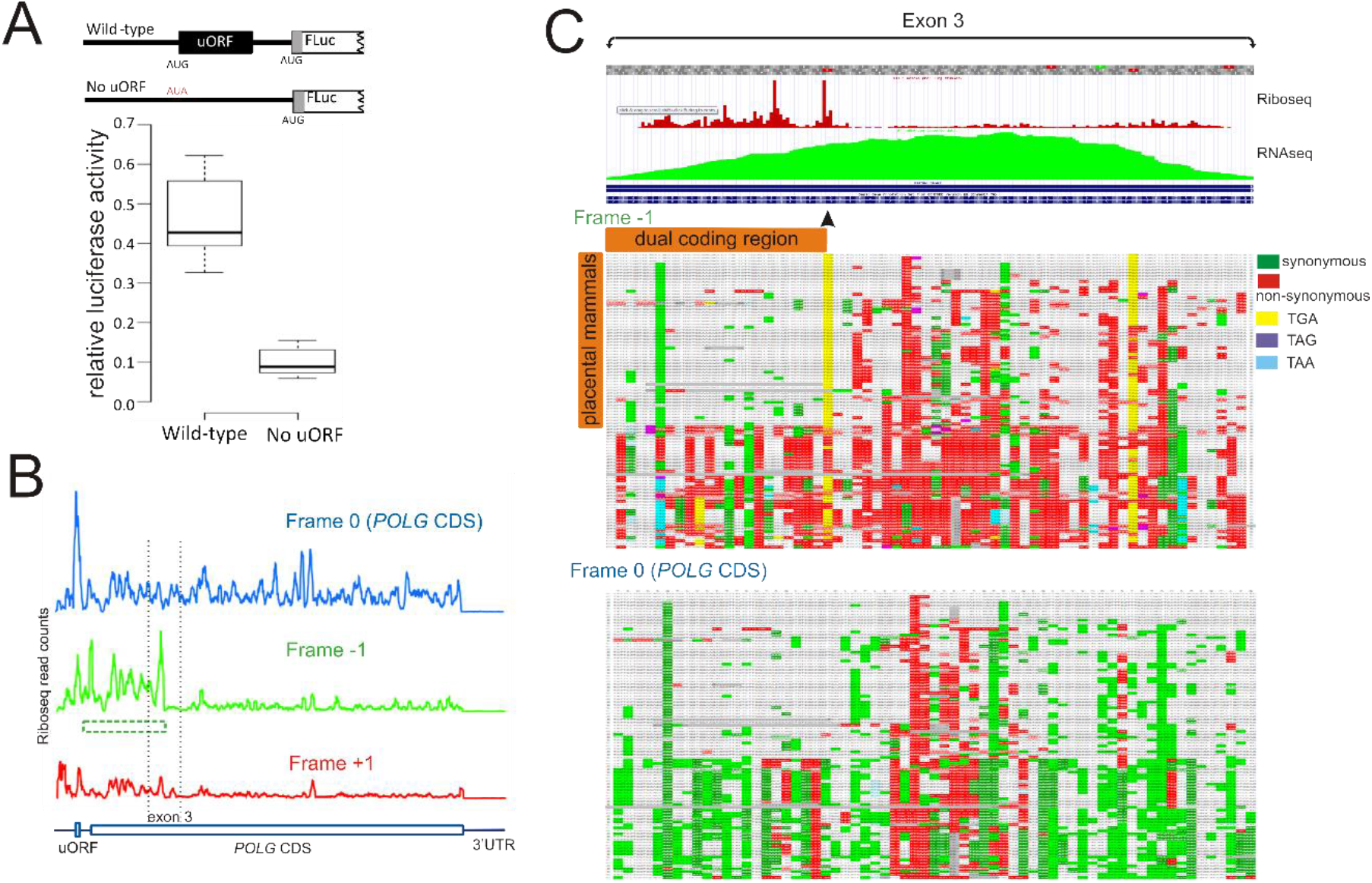
Identification of an alternative translated ORF in *POLG*. **A.** Schematic representation of reporter constructs bearing full length *POLG* 5’leaders (with and without the uORF AUG codon), its AUG start codon plus 33 nt fused to firefly luciferase (in frame with the *POLG* AUG), and relative luciferase activities of corresponding constructs transfected into HEK293T cells. **B.** Riboseq aligned to each reading frame for the *POLG* mRNA generated in the Trips-Viz browser. The region in the −1 frame which has high Riboseq density is depicted by a green dotted box. **C.** GWIPS-Viz tracks for Riboseq (red) and RNAseq (green) global aggregates for exon 3 of *POLG* (top). Lower panels represent CodAlignView alignment of 100 vertebrate genomes (hg38/100) for the −1 frame (middle) and frame 0 (bottom).

Since there are no AUG triplets that could initiate translation of the −1 frame ORF we searched for conserved near-cognate initiation codons and, in exon 2, identified a CUG triplet located 52 nt upstream of the *POLG* CDS start codon (Fig. 2A). To test whether this CUG codon can initiate translation in the alternative reading frame, we fused the 5’ leader of *POLG* to Fluc in the −1 frame (Fig. 2B). We observed robust −1 frame translation that increased by ~30% when translation of the uORF was abolished (Fig. 2B, constructs 1 and 2). Thus, translation of the uORF decreases −1 frame translation and increases 0 frame (POLG) translation (Fig. 1A), which is consistent with preferential translation reinitiation in the 0 frame after uORF translation (Fig. 1A and Fig. 2B). Replacement of the predicted CUG start codon with a non-initiating CUA completely abolished −1 frame translation, strongly suggesting that this CUG triplet is the only −1 frame initiation codon (Fig. 2B, construct 3). We termed this long ORF (260 codons in humans), which extensively overlaps with the annotated *POLG* reading frame and starts at CUG, as *POLGARF* (*<u>POLG</u> <u>A</u>lternative <u>R</u>eading <u>F</u>rame*).

**Figure 2.**
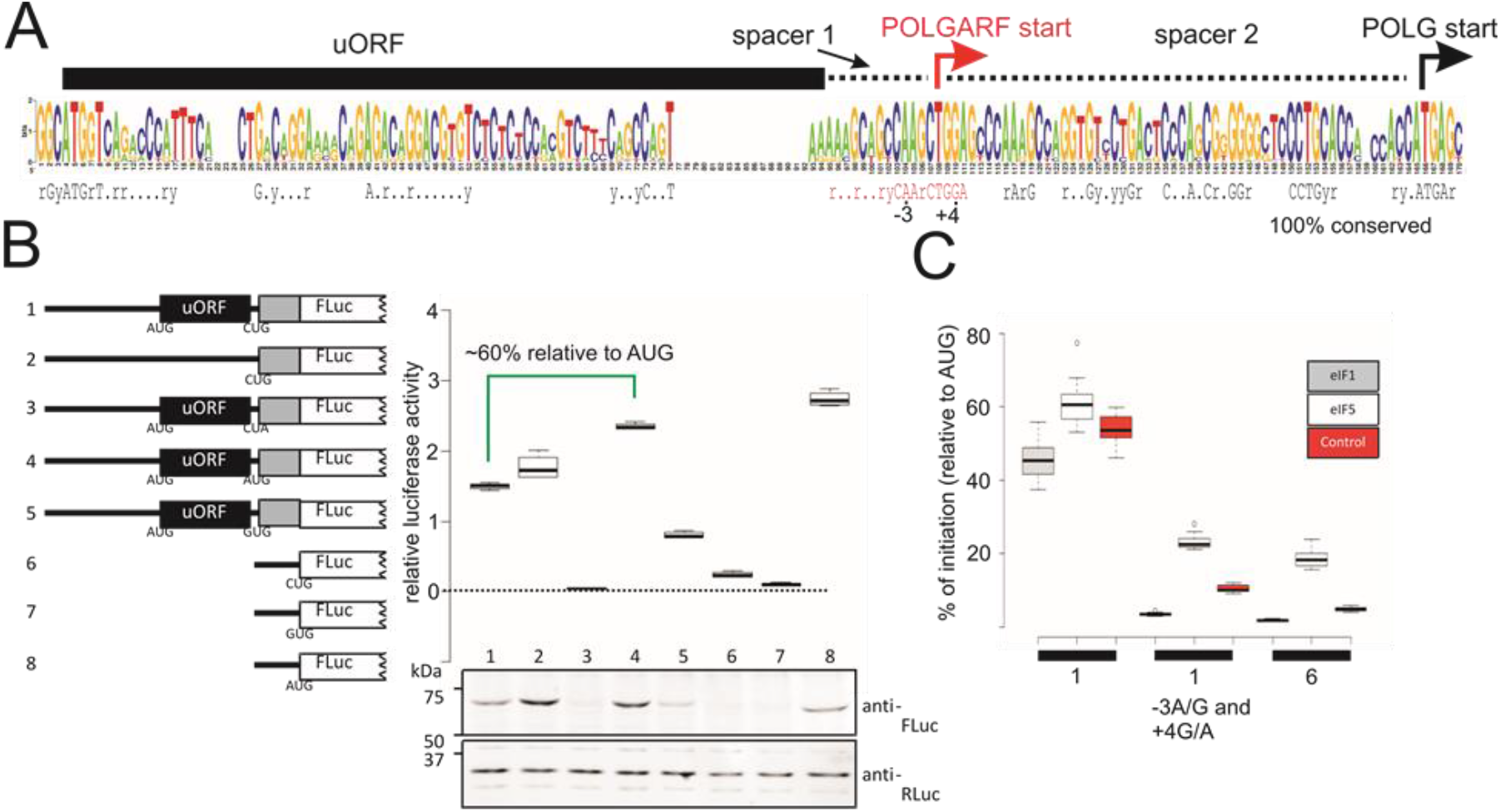
*POLGARF* translation from a CUG codon. **A.** Sequence logo of multiple alignments of a *POLG* 5’leader fragment generated for 85 placental mammals. Sequence below represents nucleotide positions that are universally conserved. The *POLGARF* CUG codon and surrounding nucleotides are highlighted in red. R and Y stands for purines and pyrimidines, respectively. **B.** Schematic representation of reporter constructs tested and results of transfection in HEK293T cells. The lower panel represents the results of Western blotting against Firefly Luciferase and control Renilla luciferase (co-transfected with constructs tested). **C.** Luciferase reporter assay of selected constructs co-transfected with plasmids expressing EIF1 or EIF5. 1 represents constructs derived from construct 1 in B. 6 = construct 6 from B.

### The POLGARF CUG acts as a highly efficient initiation codon

Notably, in reporter constructs, initiation at the *POLGARF* CUG is ~60% as efficient as an AUG in the same position (Fig 2B, constructs 1 and 4), which is markedly higher than most of the values reported for CUG initiation (5-10% efficiency, constructs 6 and 8) (25). The exceptionally high *POLGARF* CUG initiation efficiency was further explored by testing a series of 5’ - and 3’ - truncated constructs. We found that almost the entire 5’ leader is dispensable for efficient −1 frame initiation (Fig. S1). It has been reported that initiation at near cognate codons may be more dependent on local context than initiation at AUG codons (26). Therefore, it seems highly probable that such efficient *POLGARF* initiation is heavily reliant on its surrounding nucleotide context. Particularly important is a G at the +4 position (where the first nucleotide of the start codon is +1) which has been recently shown to be critical for near-cognate initiation efficiency (26). In the context of the *POLGARF* CUG, the −3A and +4G are evolutionary conserved (Fig. 2A). To check whether this purine nucleotide signature makes the observed CUG initiation so remarkably efficient, we exchanged −3A to G and +4G to A in CUG- and AUG-initiating reporters. This exchange reduced translation initiated at the AUG codon by 30% (Fig. S2), but CUG-initiated translation dropped by 85%, almost down to the levels expected for a “standard” CUG driven reporter (Fig. 2C).

Since all near-cognate initiation codons are expected to be suboptimal, it seemed likely that the *POLGARF* CUG initiation should be sensitive to the levels of initiation factors responsible for the stringency of sub-optimal start codon selection, i.e. EIF1 and EIF5 (27, 28). To test this, we co-expressed *POLG* 5’ leader reporters with an excess of either EIF1 or EIF5. Surprisingly, we saw almost no effect of EIF1 or EIF5 overexpression on *POLGARF* CUG-initiation: Thus in its natural nucleotide context the *POLGARF* CUG codon behaves as a canonical AUG codon in optimal Kozak context which is also unresponsive to EIF1 and EIF5 overexpression (29). However, the exchange of −3G and +4A completely abolishes its independence from EIF1 and EIF5 levels. (Fig. 2C). Thus, the nucleotide context of the CUG start codon rather than its identity is the major determinant of POLGARF initiation efficiency. This context is critically dependent on the combination of −3A and +4G and is also regulated by more distal nucleotides (Fig. S3).

What is the role of the short uORF upstream of *POLGARF* CUG? It is likely that ribosomes access the *POLGARF* CUG either by leaky scanning past the uORF AUG and/or by reinitiation of ribosomes that have translated the uORF. To determine whether leaky scanning is important for *POLGARF* CUG-initiation we tested reporters in which reinitiation on CUG is prevented by extending the uORF from 23 to 71 codons. Compared to wild-type constructs, CUG initiation is reduced by >50% suggesting that leaky scanning plays an important role in *POLGARF* CUG initiation (Fig. S4). In accordance with this, reducing leaky scanning by inserting a second in-frame AUG within the extended uORF almost completely abolishes CUG initiation. Confirmation that reinitiation plays an equally important role in CUG initiation is observed in constructs in which leaky scanning is abolished by insertion of an in-frame AUG but reinitiation is still possible. Here we still observe CUG initiation that is approximately half of wild-type CUG initiation. Furthermore, reducing uORF length is thought to be more permissive for reinitiation and in agreement with this we observe an approximately 10% increase in CUG initiation by decreasing the uORF from 23 to 5 codons. In conclusion, both leaky scanning and reinitiation at the uORF appear to play an equally important role in POLGARF CUG-initiation (Fig. S4).

### Being originated through MIR transposition, *POLGARF* is conserved in placental mammals

To explore whether *POLGARF* encodes a functional protein we carried out phylogenetic analysis of 100 vertebrate genomes (30). The *POLGARF* ORF as well as its CUG start codon are conserved in placental mammals, and PhyloCSF analysis (31) reveals strong purifying selection acting on the evolution of the *POLGARF* protein coding sequence (Fig. 3A).

**Figure 3.**
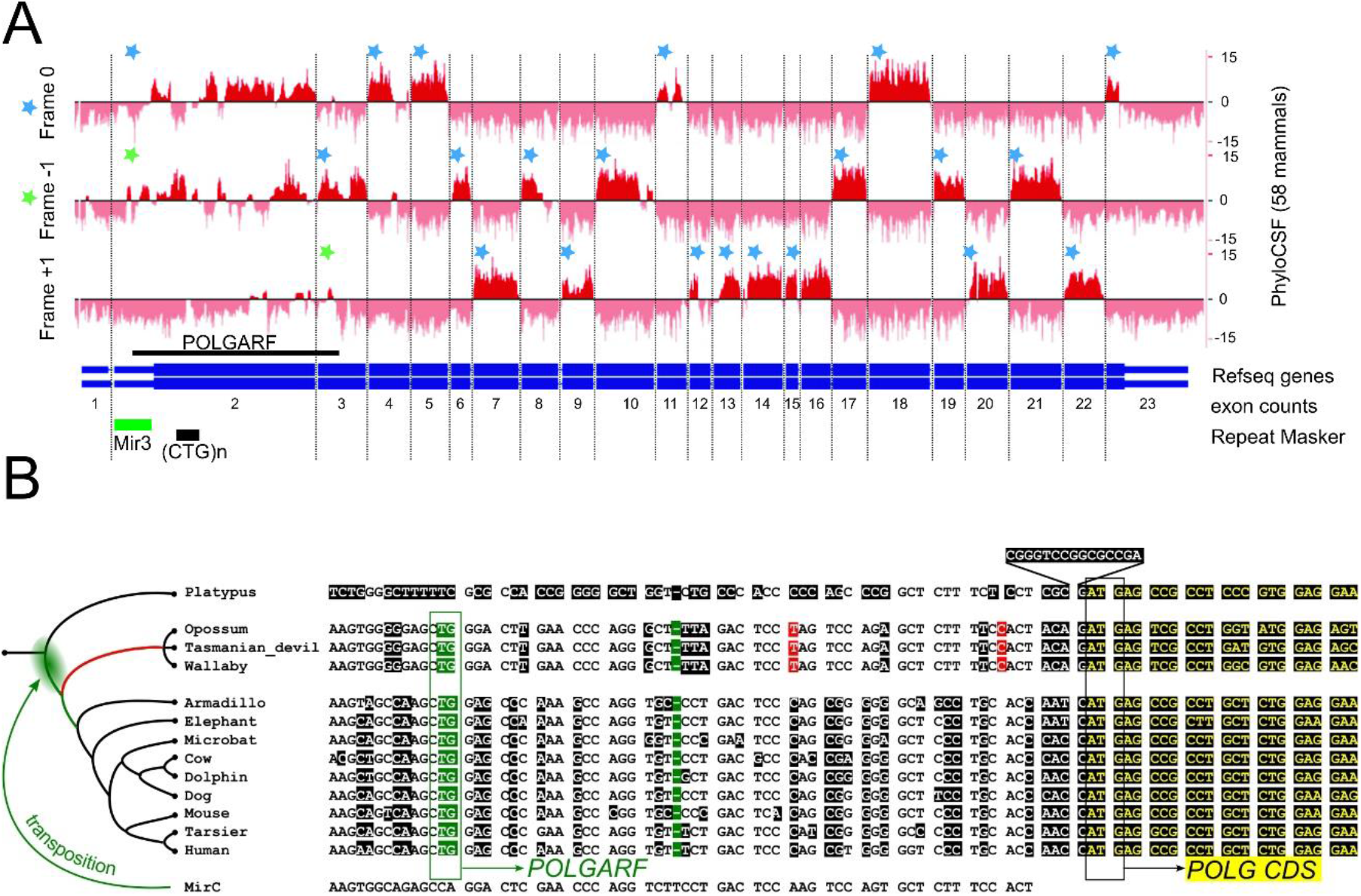
Conservation and evolution of *POLGARF*. **A.** PhyloCSF tracks for the *POLG* gene. *POLGARF* is indicated with a thick black line above Refseq annotation. RepeatMasker tracks for *POLG* exons are below ((CTG)n corresponds to a poly-glutamine track in *POLG* for minus strand). Reading frames for *POLG* and *POLGARF* ORFs are indicated with blue and green stars respectively. **B.**Possible origin of *POLGARF*. A genomic alignment is shown for representative mammalian species. Dfam/Repbase MirC consensus is given below. Nucleotides that differ from MirC sequence are highlighted: common among marsupials and placental mammals are in green; marsupial variants incompatible with *POLGARF* expression are in red; the rest are in black. *POLG* CDS sequence is in yellow. The timing of MirC sequence transposition is shown in the tree as a green cloud, the likely timing of mutations that led to the highlighted variants (in green and red) are indicated in the tree with the same colours. The initiation starts for POLG and POLGARF are indicated with rectangles.

RepeatMasker (http://repeatmasker.org) identified an MIR (mammalian-wide interspersed repeat) that overlaps with the CUG codon and the short uORF (Fig. 3A). Detailed analysis of vertebrate alignments suggests that *POLGARF* originated by an MIR transposition before adaptive radiation of placental mammals (Fig. 3B and discussion below).

### Detection of endogenous POLGARF protein

Alternative reading frame translation does not necessarily mean the existence of a stable protein product. To search for endogenous POLGARF peptides, we applied the post acquisition targeted search (PATS) technique (32) to a BioPlex interactome dataset containing AP-MS results for ~6000 protein baits overexpressed in HEK293 cells (33). The PATS algorithm predicted POLGARF peptides in >90 protein baits, among which TRIP13, CAMK2D, NPM2, HAVCR2, CLEC3A and CHCHD10 pull down datasets were predicted to have 2 or more POLGARF originated peptides (Table S1).

Direct .raw data analysis for the latter baits indeed identified four POLGARF tryptic peptides; three of these are not found within nr protein database by BLAST (34) and are unique for POLGARF. Since this is the first ever observation of POLGARF protein product, we confirmed the fidelity of peptide identification by comparing MS2 spectra from BioPlex .raw files with our LC-MS/MS data produced with overexpressed POLGARF (Fig. S5). The pattern of tryptic peptide coverage and MS2 spectra were highly similar between the two datasets. Together these data unambiguously identify endogenous POLGARF protein in HEK293 cells from the BioPlex data.

Notably, the abundance of POLGARF in cells seems to be very low, as we were unable to detect POLGARF originated peptides in either in-house generated proteomics data for HEK293 cells without POLGARF overexpression nor in deep proteome datasets from (35) for HEK293, HeLa, colon, liver, HCT116 and prostate cells, while in the same datasets several POLGARF originated peptides were detected in MCF7, SHSY, A549 cells as well as in Jurkat cells in the dataset from (36)) and in immune cells in the dataset from (37). All peptide identifications are isolated and with very low intensities. Considering the high efficiency of POLGARF translation from *POLG* mRNA observed with RiboSeq and reporter assays, it seems likely that endogenous POLGARF is either compartmentalised within subcellular locations that are insoluble under regular cell lysis protocols, or is secreted from the cell or else unstable.

### POLGARF protein interacts with C1QBP and redirects it to nucleoli

In order to shed light on POLGARF function, we searched for its interacting partners using a GST-POLGARF fusion protein overexpressed in Expi293F cells. Pull down assays detected a 32 kDa major protein identified as C1QBP, a.k.a. P32 (Fig. 4A). The P32 homotrimer adopts a doughnut-shaped quaternary structure with asymmetric charge distribution on its surface. P32 is involved in a wide range of intracellular and extracellular activities. However, directed by a mitochondrial targeting sequence (MTS), it predominantly localizes in the mitochondrial matrix (38). In mitochondria, P32 is thought to control the translation of mitochondrially encoded proteins, either directly or by affecting mitochondrial ribosome biogenesis (39, 40). We confirmed the specificity of the POLGARF/P32 interaction using SNAP-tag pull down assays: SNAP-POLGARF and SNAP-P32 fusion proteins efficiently pulled down P32 and POLGARF, respectively (Fig. 4B). Next, we found that SNAP-POLGARF, when overexpressed in HEK293T cells, accumulates in nucleoli, where it colocalizes with fibrillarin, one of the major nucleolar components (Fig. 4C), and does not colocalize with SC35, a nuclear speckle marker (Fig. S6). Nucleolar localization of POLGARF is dependent on amino acids located within its N-terminal half (Fig. S7). In nucleoli, POLGARF localizes to areas of active rRNA production, enriched with RNA polymerase I (Fig. 4C). SNAP-P32 alone was not observed in the nucleoli, however, when co-expressed with CLIP-POLGARF, SNAP-P32 showed clear nucleolar localization (Fig. 4D). To determine whether it is full-length, or mature P32 (without MTS) that accumulates in the nucleoli in a POLGARF-dependent manner, we carried out subcellular fractionation of cells overexpressing P32 with or without POLGARF (Fig. 4E). Mass spectrometry analysis of nucleolar P32 demonstrated that it retained the MTS (Fig. S8). This suggests that its interaction with POLGARF prevents P32 maturation, redirects P32 from the mitochondria to the nucleoli and thus may affect P32 functions.

**Figure 4.**
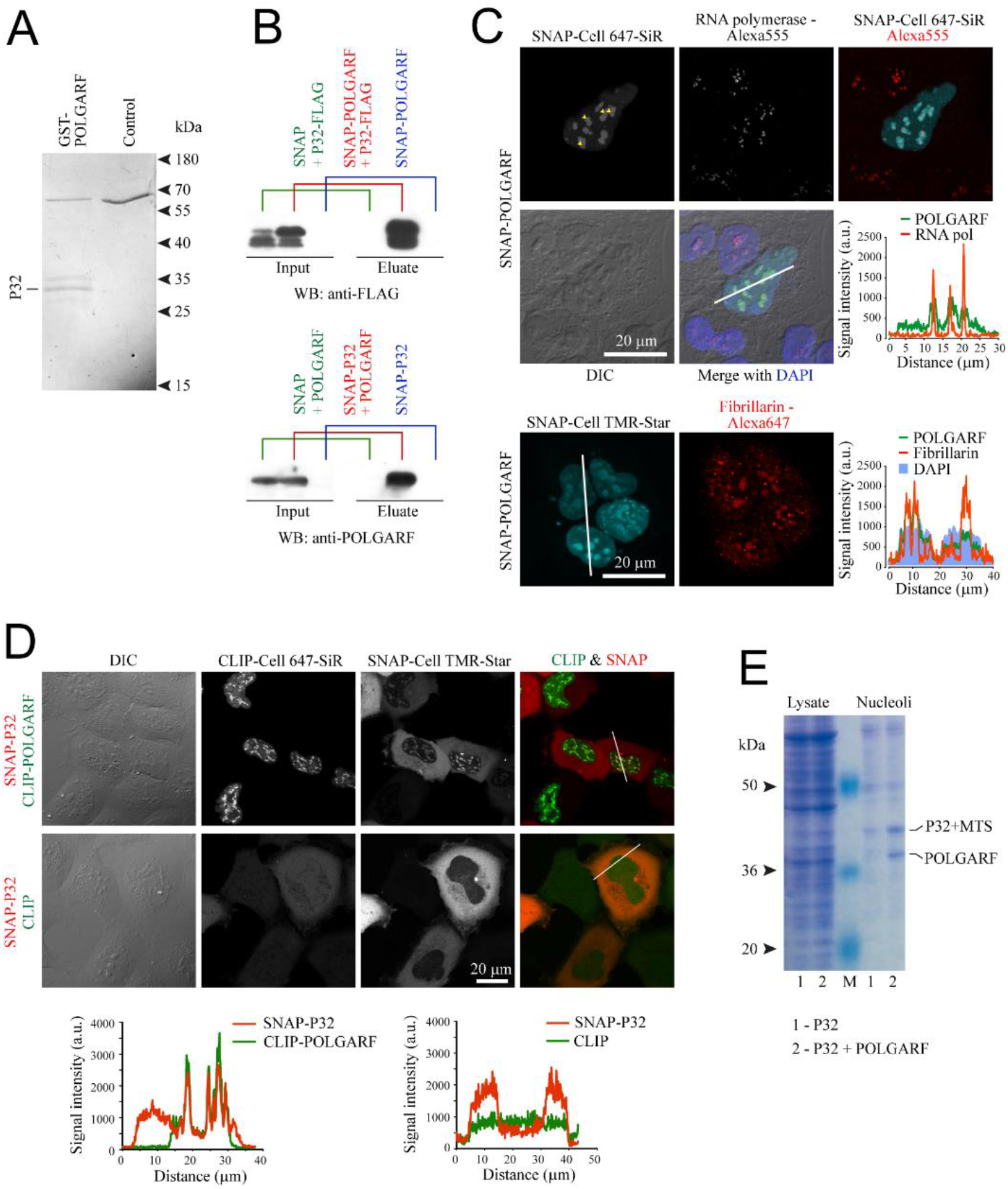
POLGARF interaction, localization and functional analysis. **A.** Coomassie blue stained SDS PAGE of a GST-POLGARF pull down assay. The major protein bands were excised and subject to mass spectrometry analysis. **B.** SNAP pull down assay from cells transfected with indicated constructs. Anti-FLAG and anti-POLGARF antibodies (see also Fig. S9) were used for Western blotting analysis. **C** and **D.** Confocal imaging of cells transfected with the indicated constructs and stained with corresponding SNAP & CLIP dyes and antibodies. Line profile analysis across representative cells shows localization of fluorescent signals. In C, yellow arrowheads show localization of RNA polymerase I (unstained spots), surrounded by POLGARF. **E**. Coomassie blue stained SDS-PAGE with total cytoplasmic lysate and with nucleolus-enriched fractions of cells transfected with the indicated constructs. Protein bands corresponding to P32 and POLGARF were excised and subject to mass spectrometry.

### POLGARF C-terminal fragment, POLGARFin, is secreted from cells upon serum stimulation

To investigate the kinetics of POLGARF accumulation, we fused POLGARF with HiBiT, an 11 amino acid peptide which can complement a truncated Nanoluciferase fragment (LgBiT) to regain full activity (41). After transfection of POLGARF-HiBit we detected a progressive increase in luciferase activity in Hek293T cell lysates. Interestingly, we also detected luciferase activity in the conditioned media (Fig. S9). Extracellular HiBiT-containing protein was purified from the conditioned media with an engineered SNAP-LgBiT protein and SNAP magnetic beads. Upon fractionation by SDS-PAGE intracellular POLGARF-HiBiT migrates at 35 kDa as expected (Fig 5A), whereas an extracellular HiBiT fusion protein migrates at ~17 kDa (Fig 5B). LC-MS analysis of the extracellular HiBit fusion identified it as a heterogenenous population of C-terminal POLGARF fragments mainly produced by cleavages around positions 138-140 and 150 with a minor population sprinkle both longer and shorter proteoforms around the major cleavage positions (Fig. 5C). We called these fragments POLGARFin. Notably, POLGARFin fragments were not observed in cell lysates representing soluble cytosolic fractions, nor could we detect full length POLGARF in the media (Fig 5A). It seems unlikely that POLGARFin-HiBiT is released into the media from dead cells. The most probable explanation is that POLGARFin is secreted immediately after intracellular POLGARF cleavage. Alternatively, the full-length protein can be secreted and immediately cleaved outside of cells.

**Figure 5.**
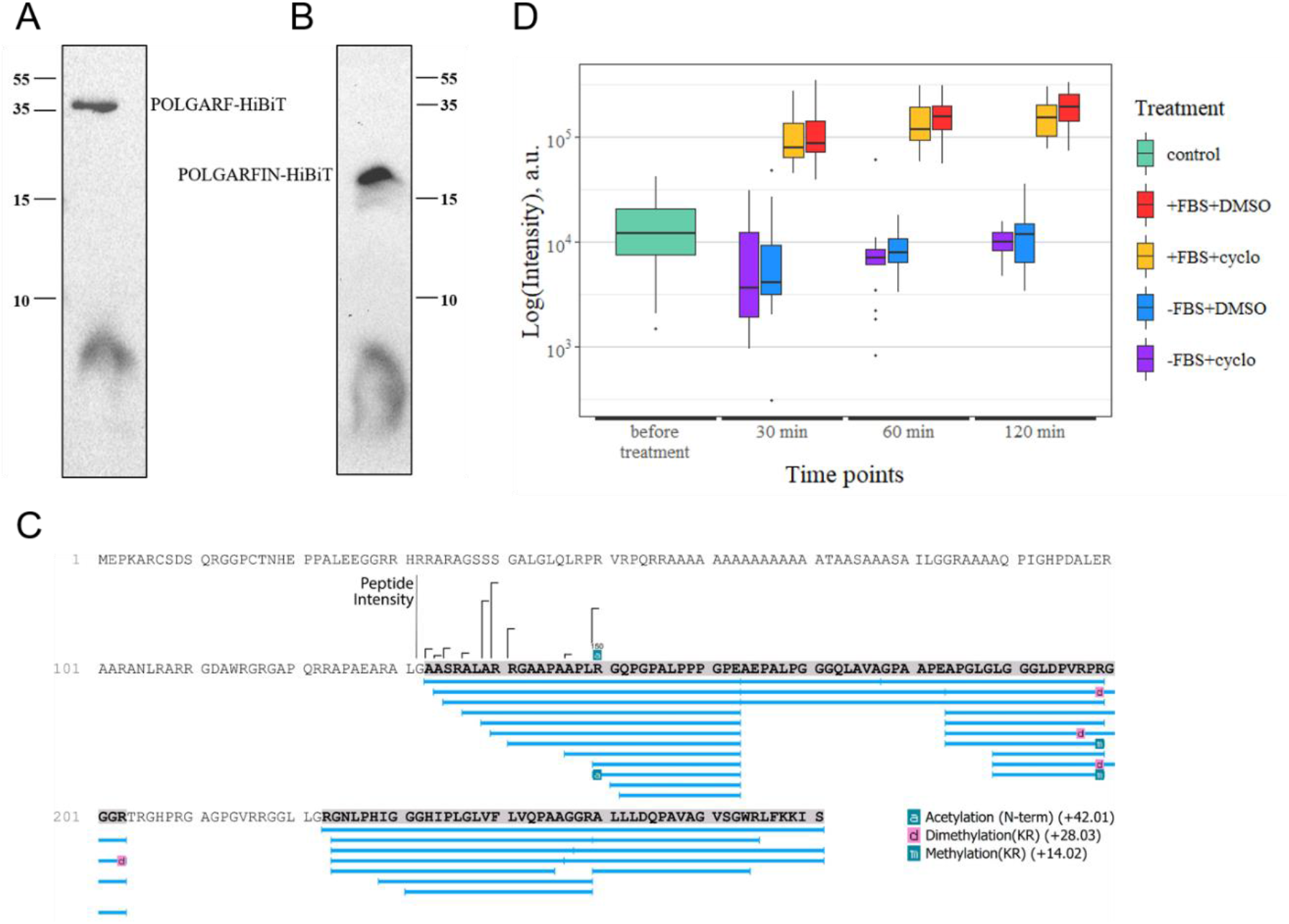
POLGARF cleavage and secretion of its C-terminal fragment upon serum stimulation. **A.** HiBiT-blotting of cell lysate after expression of POLGARF-HiBiT. **B.** HiBiT-blotting of secreted HiBiT containing polypeptide purified from conditioned media from cells expressing POLGARF-HiBiT. **C**. LC-MS sequencing of secreted POLGARFin-HiBiT. Peptide peak areas are shown for the sequences at the ragged N-term of POLGARFin. **D.** Measurement of HiBiT activity in the conditioned media from POLGARF-HiBiT transfected Hek293T cells. 24h after transfection, the media was replaced with fresh DMEM +/−10% FBS and +/-cycloheximide, after indicated time points the media aliquotes were assayed with Nano-Glo Nano-Glo HiBiT luciferase assay (n=24).

To investigate whether this secretion is regulated, we subjected POLGARF-HiBiT overexpressed cells to various treatments. It appeared that POLGARFin secretion is upregulated by serum addition: when the media of transfected cells is supplemented with fresh media containing 10% FBS, rapid accumulation of POLGARFin is observed within 30 minutes after stimulation with no further increase over time (Fig 5C). In contrast, addition of serum free media does not result in POLGARFin secretion. POLGARFin secretion in response to FBS-containing media does not depend on *de novo* protein synthesis, as supplementation of cells with serum-rich media infused with cycloheximide does not prevent POLGARFin extracellular accumulation. Collectively, these data suggest that POLGARF is processed into actively secreted POLGARFin, which may be implicated in extracellular signalling events.

## Discussion

The results presented above provide strong evidence for the unusual existence of a novel functional protein encoded within the *POLG* mRNA. When two different proteins are translated from a single mRNA, the efficiency of initiation at each start codon could set the ratio of product steady state levels assuming each protein has similar stability. However, this ratio may vary during conditions in which initiation efficiency is altered. Eukaryotes have developed elaborate mechanisms for the recognition of the correct initiation codon and the levels of certain initiation factors can regulate the fidelity of initiation, especially on sub-optimal (near-cognate and AUG in poor context) start codons (42). While elevated levels of EIF1 can increase the stringency of start codon selection, elevated levels of EIF5 have the opposite effect. Here we show that overexpression of either EIF1 or EIF5 had minimal effect on CUG-initiated POLGARF (Fig. 2C) suggesting that this highly efficient CUG initiation is refractory to normal stringency controls.

Our analysis of the role of the conserved uORF in POLGARF initiation reveals that both leaky scanning and reinitiating ribosomes can equally start translation at the POLGARF CUG (Fig. S4). However, since preventing translation of the uORF (therefore no reinitiation) results in predominantly CUG initiation then it follows that under normal conditions most of the ribosomes initiating the POLG ORF are reinitiating ribosomes. This raises the intriguing possibility that the ratio of POLG to POLGARF may be regulated by stress conditions.

Although the contribution of SINE transposons to eukaryotic gene evolution, including their exonization (43), is well known, to our knowledge this is the first example of a retrotransposon creating a dual coding gene. How did POLGARF evolve and why did it subsequently become fixed in placental mammals? All mammalian sequences with the exception of platypus share significant similarity with MIR sequences as determined with Dfam search (44). Therefore, it is likely that MIR insertion occurred after Monotremata diverged from the common ancestor of Marsupials and Placentals. We found that POLG sequences of many vertebrate species (including platypus) lack stop codons in one of the alternative frames in the first coding exon, thus, acquisition of an in-frame start codon could lead to the expression of the alternative frames. However, MIR did not contain a suitable start codon. Two subsequent mutations had to occur to enable POLGARF expression: a substitution of CCA with CTG and a single nucleotide deletion downstream that was necessary to place the CTG in-frame with the alternative ORF. Interestingly, these variants are common for both marsupials and placental mammals (shown in green in Fig 3B), suggesting that proto-POLGARF existed in their common ancestor. However, marsupials have two variants (in red in the region shown in Fig 3B) and several stop codons in the POLGARF frame further downstream, suggesting that POLGARF was subsequently lost in marsupials, while in the common ancestor of placental mammals it acquired a functional role and became fixed in subsequent lineages.

What may be the functional role of POLGARF? The polypeptide can be tentatively divided into four parts; notably, the 64 amino acid long C-terminus is the most conserved region with 22 invariant amino acids (Fig. S10). We failed to find any significant similarity between POLGARF and other known or predicted proteins or any similarity with known structural motifs. It seems likely that POLGARF is an intrinsically disordered protein (IDP) with a remarkably high isoelectric point (pI =12.05 for a human protein).

As a conserved IDP, POLGARF has a good potential to be an important regulatory protein. In cell signalling and regulation, IDPs emerged as parts of integrated circuits. Indeed, the capacity of IDPs to acquire numerous posttranslational modifications and change conformation in a context-dependent manner allows immense versatility of their interactomes (45, 46). The observed specific interaction of POLGARF with P32 as well as the modulatory effects of POLGARF on P32 localisation and processing exemplify the importance of POLGARF for cell functioning (Fig 4 and Fig S8). Along with P32, other putative interaction partners of POLGARF (TRIP13, CAMK2D, etc., see Table S1) can be considered as good candidates for mechanistic follow up studies.

According to RiboSeq analysis and reporter assays, the levels of POLG and POLGARF proteins should be comparable. However, according to proteomics data, in contrast to the moderately high levels of the housekeeping protein POLG, POLGARF’s concentration is extremely low. Our discovery that POLGARFin can be secreted is likely the most probable explanation for this discrepancy. We propose that endogenous POLGARF may be almost completely processed and secreted outside of cells, where it may participate in currently unknown cell-to-cell communication. We speculate that the levels of secreted POLGARFin reflects the capacity of the donor cell to have enough POLG to replicate mtDNA, as both POLG and POLGARF are encoded in the same mRNA. In contrast, cells with decreased expression of *POLG* mRNA would also produce less POLGARFin.

Finally, these findings could have profound implications for the interpretation of POLG mutations. As a previously unknown dual coding gene, *POLG* could bear hidden mutations responsible for diseases of still unknown aetiology. Among the known *POLG* mutations some do not cause changes in the amino acid composition of POLG. Such synonymous single nucleotide variations (SNVs) are often considered harmless (except for their potential effects on splicing) (47). However, in dual coding regions (such as POLG/POLGARF described in this study), synonymous SNVs in one ORF are unlikely to be synonymous in the other ORF. This emphasizes the need to consider *POLGARF* variants in future studies.

## Materials and Methods

### Cell culture

Here and elsewhere, all reagents were from Millipore-Sigma, unless stated otherwise. Human embryonic kidney HEK293T and Expi293F cells were from the American Collections of Cell Cultures (Manassas, VA) and Thermo Fisher Scientific (Rockford, Ill), respectively. HEK293T were maintained in 75 cm2 flasks (Sarstedt, Wexford, Ireland) in DMEM supplemented with 10% FBS, 2 mM L-glutamine, 100 U/ml penicillin / 100 μg/ml streptomycin (complete DMEM) with or without 10 mM HEPES (pH 7.2), in humidified atmosphere of 95% air and 5% CO_2_ at 37°C. Expi293F cells were maintained in Expi293 Expression medium (Thermo Fisher) at 8% CO_2_ and 37°C under continuous mixing at 160 RPM on an orbital shaker.

### Plasmids and constructs

To create plasmids for luciferase reporter assays, the POLG 5’ leader plus 33 nt of POLG CDS were amplified by PCR from HEK293T cDNA using sense primer 5’-GACCCAAGCTTAAGTTGCGGCTGCCAGCG and antisense primer 5’-TTATTGGATCCCCGGCCACCTTCCTCCAGAGC with flanking HindIII and BamHI restriction sites. All other variants were generated by 2-step PCR using appropriately designed mutagenic or nested deletion primers. All PCR amplicons were digested with HindIII and BamHI and ligated into HindIII and BamHI digested dual luciferase vector p2-Luc (48) to make pSV40-firefly reporter constructs indicated. All constructs were verified by sequencing. Plasmids used to overexpress deregulated eIF1 (eIF1g*), and deregulated eIF5 (eIF5-AAA), have been described previously (29). Invitrogen pcDNA3 plasmid (Thermo Fisher) was used for control transfections.

To create plasmids for full length POLGARF-FLAG overexpression, the region of the POLG mRNA corresponding to the complete 5’ leader to the POLGARF stop codon was amplified by PCR from a HEK293T cDNA library using a sense primer containing a 5’ HindIII site and an antisense primer containing FLAG in frame to POLGARF followed by an XbaI site. This amplicon was cloned between HindIII and XbaI in pcDNA3.1 (Thermo Fisher) to create pcDNA3.1 POLGARF-FLAG. In parallel, a plasmid without the full 5’leader and with a CUG to AUG mutation was created using the same reverse primer containing FLAG (pcDNA3.1 POLGARF AUG-leader).

The POLGARF and P32 coding sequences were first amplified and inserted into Invitrogen pGEX6p1 vector (Thermo Fisher) between BamH1 and NotI sites (in frame to N-terminal GST tag) to create pGEX6p1-POLGARF and pGEX6p1-P32 respectively. These constructs were used to prepare the subsequent constructs used in this study. First, the GST-POLGARF region was amplified from pGEX6p1-POLGARF using a forward primer which modifies the GST start codon to an efficient initiator in mammalian cells. This fragment had primer-incorporated XbaI and NotI sites and was inserted into a modified pcDNA3.4 vector (Thermo Fisher), which contained a custom polylinker (with XbaI and NotI restriction sites), thus yielding pcDNA3.4-GST-POLGARF.

To create constructs with SNAP and CLIP tags, the pGEX6p1-POLGARF and pGEX6p1-P32 were digested with BamH1 and NotI and cloned into pSNAPf and pCLIPf vectors from New England Biolabs (NEB, Ipswich, MA) to create pSNAPf-POLGARF, pCLIPf-POLGARF, pSNAPf-P32 and pCLIPf-P32, respectively. These constructs were further re-cloned into pcDNA3.4 (Spe1-Not1 digestion) to create pcDNA3.4-SNAP-POLGARF, pcDNA3.4-CLIP-POLGARF, pcDNA3.4-SNAP-P32 and pcDNA3.4-CLIP-P32, respectively.

To create the plasmid containing POLGARF with the C-terminal HiBiT tag, pcDNA3.4-POLGARF was modified by amplification of POLGARF with a reverse primer containing the HiBiT sequence and NotI. The resulting fragment was digested with HindIII and NotI, then again inserted in pcDNA3.4 to yield pcDNA3.4-POLGARF-HiBiT.

To create the plasmid containing SNAP-LgBiT, the gene block containing LgBiT sequence flanked by BamH1 and NotI was ordered from IDT. pcDNA3.4-SNAP-POLGARF plasmid was treated with BamH1 and NotI, and POLGARF coding fragment was exchanged with LgBiT sequence to yield pcDNA3.4. SNAP-LgBiT.

The P32 CDS with C-terminal FLAG was cloned in pcDNA3.4 to create pcDNA3.4-P32-Flag. The construct pcDNA3.1 POLGARF AUG-leader was recloned into pcDNA6 vector to create pcDNA6-POLGARF-Flag.

The pcDNA3.4-SNAP-POLGARF construct was used to generate the series of N-terminal and C-terminal deletions: ΔN1 to ΔN5 corresponds to progressive 26 aa deletions from its N-terminus, while ΔC1 to ΔC5 corresponds to progressive 26 aa deletions from its C-terminus (Fig S7). The sequences were amplified with corresponding primers bearing BamH1 and Not1 restriction sites and were cloned back into pcDNA3.4 to generate truncated fusion proteins.

### Cell Transfection

Four main transfection protocols were used in different layouts.

1. One-day Lipofectamine 2000 based protocol, in which suspended cells are added directly to the DNA complexes prepared in OptiMEM. For luciferase assays, HEK293T cells were transfected in half-area 96-well or 6-well plates (Sarstedt). For Figs. 1A, 2B and Supplementary Figs. 1-3, the following were added to each well: 25 ng of each pSV40-firefly vector, 5 ng control pSV40-Renilla vector plus 0.2 μl Lipofectamine 2000 in 25 μl OptiMEM (both from Thermo Fisher). For Fig. 2C, the following were added to each well: 50 ng of protein overexpressing vector, 5 ng pSV40-firefly vector (with initiation contexts and/or codons as indicated in the figures), 1 ng pSV40-Renilla vector and 0.2 μl Lipofectamine 2000 in 25 μl OptiMEM. The transfecting DNA complexes in each well were incubated with 3×10^4^ cells suspended in 50 μl DMEM + 10% FBS at 37°C in 5% CO_2_ for 24 hr. For Western blotting analysis of luciferase expression, 10^6^ HEK293T cells suspended in 3 ml DMEM + 10% FBS were transfected overnight with 1 μg of each indicated plasmid in 6-well plates.
2. One-day Lipofectamine 2000 based protocol, in which DNA complexes prepared in OptiMEM are added to 50-60% confluent adherent cells. For confocal imaging and Western blotting analysis of antibodies against POLGARF, HEK293T cells were seeded at 1.5×10^4^ cells /cm^2^ on glass bottom dishes (MatTek, Ashland, MA)) or 2.5×10^5^ per well of 6-well plates (Sarstedt). After 12 h incubation, plasmids encoding construct as indicated in Figures were delivered in cells by 3-4 h incubation with DNA complexes; 100 ng DNA and 0.4 μl Lipofectamine were used per 1 cm^2^. Then transfection mixture was replaced with standard medium and cells were grown for 20 h prior to lysis or SNAP/CLIP staining followed by live cell microscopy or immunostaining.
3. FuGene protocol (40-48 h). For GST- and SNAP-pull down assays, HEK293T cells were transfected with combinations of pcDNA3.4-SNAP-POLGARF, pcDNA3.4-CLIP-POLGARF, pcDNA6-POLGARF-Flag, pcDNA3.4-P32-Flag, and pcDNA3.4 encoding fluorescent protein (Fast Fluorescent Timer) (49) as a negative control (2 μg of DNA per well of 6 well plate) with FuGene transfection reagent (Promega, Madison, WI) according to manufacturer’s instructions.
4. Expifectamine 293 protocol (48 h). For subcellular fractionation, 2 × 10^8^ expi293F cells were transfected with DNA. For that, 80 μg DNA and 216 μl Expifectamine 293 (Thermo Fisher) were prepared in 4 ml OptiMEM, mixed together, incubated for 20 minutes at room temperature and added to the cells. In 24 h, enhancers 1 and 2 (400 μl /4 ml) and gentamicin (up to 10 μg/ml) were added. Cells were harvested by centrifugation (10 min, 3000 g) 48 h after transfection.

### Dual Luciferase Assay and Western analysis

Firefly and Renilla luciferase activities were determined using the Dual Luciferase Stop & Glo® System (Promega). Relative light units were measured on a Veritas Microplate Luminometer with two injectors (Turner Biosystems, Sunnyvale, CA). Transfected cells were lysed in 12.6 μl of 1 × Passive Lysis Buffer (PLB, Promega) and light emission was measured following injection of 25 μl of either Renilla or Firefly luciferase substrate. Initiation efficiencies (% initiation) were determined by calculating relative luciferase activities (Firefly/Renilla) of test constructs and dividing by relative luciferase activities from replicate wells of control ATG constructs.

For Western blotting analysis of luciferases, transfected cells were lysed in 100 μl 1 × PLB. Proteins were resolved by SDS-PAGE and transferred to Protran nitrocellulose membranes (GE Healthcare Life Sciences, Waukesha, WI), which were incubated at 4°C overnight with primary antibodies. Immunoreactive bands were detected on membranes after incubation with appropriate fluorescently labelled secondary antibodies using a LI-COR Odyssey® Infrared Imaging Scanner (LI-COR, Lincoln, NE).

For Western blotting analyses of POLGARF expression, transfected cells were washed with PBS and lysed for 20 min on ice with RIPA buffer (Thermo Fisher), containing phosphatase and protease inhibitors; complete protease inhibitor cocktail and PhosSTOP tablets were from Roche (Mannheim, Germany). After lysate clarification by centrifugation for 15 min at 14000 g and 4°C, protein concentration was measured using BCA^TM^ Protein Assay kit (Thermo Fisher) and equalised. Proteins were separated by 4-20% polyacrylamide gel electrophoresis using pre-made acrylamide gels and running buffers from GeneScript (Piscataway, NJ), transferred onto a 0.2 μm ImmobilonTM-P PVDF membrane (Sigma) using Hoefer™ TE 22 transfer system (Hoefer, Holliston, MA) and probed with antibodies against POLGARF (1:1000) and α-tubulin (1:5,000) in 5% fat-free milk in TBST (0.8% Tween-20). Immunoblots were analysed using Amersham™ ECL™ Prime Kit from GE Healthcare Life Sciences (Waukesha, WI) and the LAS-3000 Imager (Fujifilm, Japan). Quantitative image analysis was performed with ImageJ program using α-tubulin signals for normalisation.

### GST and SNAP pull down assays

For GST pull down assay, 2 × 10^8^ expi293F cells transfected with pcDNA3.4-GST-POLGARF or with pcDNA3.4-GST were harvested by centrifugation and lysed in 2 ml of PLB. The lysates were diluted with PBS to 10 ml and incubated with 200 μl of GST-Sepharose (GE Healthcare) for 1 hour on ice under agitation. GST resin was washed 3 times with 10 ml of PBS, and POLGARF bound proteins were eluted by incubation with 1 μl of Prescission Protease (GE Healthcare) in PBS at 4°C overnight. The eluted proteins were resolved by SDS-PAGE and stained with Coomassie, protein bands were excised and subjected to MALDI TOF MS/MS analysis.

For SNAP pull down assay, transfected cells were lysed in 200 μl PLB supplemented with 1 mM DTT. The lysates were mixed with 30 μl of SNAP-magnetic beads (NEB) and incubated for 1 h at 24°C on a thermomixer (900 rpm). After incubation, the beads were washed twice with 1 ml of PBS supplemented with 1 mM DTT, and bound proteins were eluted by boiling with SDS gel loading buffer. The samples were resolved on SDS page and immunoblotted with either anti-FLAG or with custom made anti-POLGARF antibodies.

### Staining of cells with probes and confocal microscopy

Transfected cells were stained with SNAP-Cell® TMR-Star or SNAP-Cell 647-SiR (NEB), both diluted 1:500 with complete DMEM, for 30 min immediately prior to live cell imaging or immunostaining. Live cell imaging was conducted on an Olympus FV1000 confocal laser scanning microscope with controlled CO_2_, humidity and temperature. Alexa-555 and SNAP-Cell® TMR-Star probes were excited at 543 nm (1-10% of laser power) with emission collected at 560-600 nm. Alexa-647 and SNAP-Cell 647-SiR dyes were excited at 633 nm (2-10% of laser power); emission was collected at 640-700 nm. Separate spectral signals were collected with 0.5 μm steps (80-120μm aperture) in sequential laser mode with emission gates adjusted to avoid spectral overlap. Fluorescence and differential interference contrast (DIC) images were collected using oil immersion UPLSAPO 60x/1.35 Super Apochromat objective. Analysis was performed using FV1000 Viewer software (Olympus) and Microsoft Excel.

### Immunofluorescence analysis

Immunostaining was performed according to a standard procedure. Briefly, cells were washed with Dulbecco’s PBS (supplemented with Ca^2+^ and Mg^2+^) fixed with 3.7% paraformaldehyde in PBS, quenched with 50 mM NH_4_Cl, permeabilised with 0.25% TX100, blocked with 5% BSA in TBST and incubated with primary and secondary antibodies for 1 h (as indicated in Figures, Supplemental Table 3). Standard TBST was used for cell washes. Commercial primary antibodies were diluted in blocking solution 1:500-1:800; POLGARF antibodies − 1:300; secondary antibodies −1:1000. When required, cell nuclei were stained with DAPI. Cells were left in PBS for confocal imaging performed within 4 h as described above for live cells. DAPI was excited at 405 nm with emission collected at 420-460 nm.

### Subcellular Fractionation

Nucleolar fraction was prepared from suspension expi293F cells according to the protocol described by Lamond and co-authors (50) with modifications. 2 × 10^8^ cells were harvested by centrifugation 48 h after transfection. Cell pellets were re-suspended in 2 ml of hypotonic buffer A (10 mM HEPES (pH 7.9), 10 mM KCl, 1.5 mM MgCl_2_, 0.5 mM DTT and Complete Protease Inhibitor tablet), incubated on ice for 15 min, homogenized 25 times in a Dounce tissue homogenizer (using a tight pestle ‘B’) and centrifuged at 1000 g for 5 min. Pellets were re-suspended in buffer A, incubated on ice for 10 min, then homogenization followed by centrifugation was repeated in order to obtain purer nuclear pellet. Then the pellet was re-suspended in 3 ml of S1 solution (0.25 M Sucrose, 10 mM MgCl_2_ and Complete Protease Inhibitor tablet), layered over 3 ml of S2 solution (0.35 M Sucrose, 0.5 mM MgCl_2_ and Complete Protease inhibitor tablet), and centrifuged at 1430 g for 5 min. Purified nuclear pellets re-suspended in 3 ml of S2 solution were sonicated 5 × 10 seconds using SONICS Vibra cell (Sonics & Materials inc, Newton, CT) (50% of max power). Sonicated sample was layered over 3 ml of S3 solution (0.88 M Sucrose, 0.5 mM MgCl_2_ and Complete Protease Inhibitor tablet) and centrifuged at 3000 g for 10 min. The resulting pellet contained the nucleolar enriched fraction.

### MALDI TOF analysis

Protein spots were excised from the gel and digested with trypsin. The spots (3-4 mm^3^) were destained with 50 mM ammonium bicarbonate 40% acetonitrile then dehydrated with 100 μL acetonitrile (ACN) and rehydrated with 5 μL of digestion solution containing 20 mM ammonium bicarbonate and 15 ng/μL sequencing grade trypsin (Promega). Digestion was carried out at 37°C for 6 h. Peptides were extracted with 10 μL of 0.5% trifluoroacetic acid (TFA) solution. 1 μL of extract was mixed with 0.5 μL of 2,5-dihydroxybenzoic acid saturated solution in 20% ACN and 0.5% TFA on the stainless steel MALDI sample target plate. Mass spectra were recorded on an UltrafleXtreme MALDI-TOF/TOF mass spectrometer (Bruker Daltonics, Germany) equipped with an Nd laser (354 nm). The MH^+^ molecular ions were detected in reflector-mode in the mass range 600-6000 m/z. The accuracy of mass peak measurement was within 30 ppm. Peptide fragmentation spectra were obtained in LIFT mode with the accuracy of daughter ions measurement within 1 a.m.u. range. Mass-spectra were processed in FlexAnalysis 3.3 software (Bruker Daltonics) and analysed manually. Protein identification was performed by Mascot software (Matrix Science, London, UK) against UniProt reference proteome (UP000005640) database. Partial oxidation of methionine and propionamidomethylation of cysteine residues was assumed for peptide mass fingerprint searches. Protein scores greater than 70 were considered as significant (p< 0.05).

### Identification of POLGARF in proteomics data

Using the recently developed Post-Acquisition Targeted Searches technique (32), we submitted the 8 longest predicted tryptic peptides from POLGARF for a targeted search in the pre-analysed HEK293 interactome dataset (33) through http://www.pepchem.org/PATS. The search yielded a total of 118 .raw file hits for 94 protein baits (Table S1). For the baits with >2 predicted peptide identifications, all .raw data were downloaded from BioPlex2.0 website (http://bioplex.hms.harvard.edu) for in-house database search analysis.

POLGARF predicted tryptic peptides were searched in PeptideAtlas (51). One of the peptides AAAAQPIGHPDALER turned out to be known and identified in several proteome datasets mostly connected with immune cells. In particular this peptide was identified in Jurkat dataset from (36) and in immune cell analysis from (37). For some other cell lines, including HEK293 cells, the dataset was taken from (Bekker-Jensen et al., 2017). Corresponding datasets were downloaded from ProteomeExchange for in-house database search analysis. For the list of all publicly available datasets downloaded and reanalysed against POLGARF-containing protein database see Supplementary Table 4.

To obtain reference peptide spectra, POLGARF was overexpressed in HEK293T and HEK 293F cells. 293T cells were transfected in 12 well plates with 1 ug of pcDNA3.4 POLGARF or pcDNA3.4 Timer (as a negative control) with Fugene transfection reagent (Promega) according to manufacturer’s instructions. After 48hr the cells were washed with PBS and lysed in a sonication bath in 150 ul of 1x Passive Lysis Buffer (Promega) supplemented with Protease Inhibitor cocktail (Roche). Expi293F cells were transfected with pcDNA3.4 POLGARF or pcDNA3.4 (40 ug of DNA per 108 of cells) with Expifectamine 293 (Thermo Fisher). After 48hr cells were washed with PBS and lysed in a sonication bath in 1x Passive Lysis Buffer (Promega) supplemented with Protease Inhibitor cocktail (Roche). The lysates were heated for 10 min at 90°C. Protein material was purified by MeOH-chloroform precipitation (52) and resuspended in 1% SDC, 100 mM Tris-HCl, pH 8.5 buffer. Protein concentration was measured by microBCA assay and 50 μg of protein was readjusted to the concentration 1 mg/ml with the 1% SDC, 100 mM Tris-HCl, pH 8.5 buffer, cysteine disulphide bonds were reduced and alkylated by adding TCEP and CAA to 10 and 20 mM correspondingly and heating at 85°C for 10 min. For protein digestion, trypsin (Promega, USA) was added at 1/100 w/w ratio twice with the first digestion for 2 h and the second digestion overnight at 37°C. Digestion was stopped by adding TFA to 1%. Precipitated SDC was removed by centrifugation. The samples were directly analysed by LC-MS without SPE.

LC-MS analysis was done on an Ultimate 3000 RSLCnano HPLC system connected to a QExactive Plus mass spectrometer (Thermo Fisher). Samples were loaded on a home-packed 2 cm × 100 μm precolumn packed with Inertsil ODS3 3 μm sorbent (GLSciences, Japan) in loading buffer (2% ACN, 98% H_2_O, 0.1% TFA) at 10 μl/min flow rate and separated on a 50 cm × 100 μm analytical column home-packed (53) with Reprosil Pur C18 AQ 1.9 μm sorbent (Dr. Maisch HPLC GmbH, Germany) in 360 μm OD 100 μm ID polyimide coated fused-silica capillary with a laser-pulled emitter prepared on P2000 laser puller (Sutter, USA). Peptide separation was carried out with a 2 h gradient of 80% ACN, 19.9% H_2_O, 0.1% FA (buffer B) in 99.9% H_2_O, 0.1% FA (buffer A) at RT.

MS data were collected in DDA mode with the following parameters: MS1 resolution 70K, 3e6 AGC target with 30 msec IT, 350-2000 m/z scan range; MS2 resolution 17.5K, 1e5 AGC target with 50 msec IT, 10 dependent MS2 scans, 1.4 m/z isolation window with 0.2 m/z offset, NCE 27; min AGC 8e3, charge exclusion unassigned, 1 and >7, preferred peptide match with isotope exclusion and 40 sec dynamic exclusion.

Raw files from BioPlex, from ProteomeExchange and the in-house generated LC-MS data were subjected to protein identification in Peaks Studio X (Bioinformatic Solution Inc., Waterloo, CA) against the UniProt SwissProt Homo Sapiens database containing both canonical and isoform proteoforms (version from 2019.08.26) with a manually attached POLGARF sequence. Search parameters included trypsin D|P digestion with max 3 miscleavages, precursor mass correction, 10 ppm and 0.05 Da error tolerance for precursor and fragment ions respectively, oxidation (M) and deamidation (NQ) as variable modifications (max number of variable modification per peptide = 5) and carbamidomethylation (C) as a fixed modification, Decoy-Fusion FDR estimation. Identification results were filtered by 0.1% PSM FDR and 1 unique peptide per group with the final protein FDR <1%.

### Induction of POLGARFin secretion

In initial experiments Hek293T cells were transfected in three 4-well plates with POLGARF-HiBiT (FuGene protocol). After every 24 hr a plate was taken out, medium and cells were harvested, and luminescence in cell lysates and in the media was measured using Nano-Glo HiBiT luciferase assay. This experiment was repeated 5 times.

To investigate the conditions that mediate POLGARFin secretion, Hek293T cells were transfected with POLGARF-HibiT in 48-well plate (FuGene protocol). After 24 hr post transfection, medium was substituted for a fresh DMEM with or without 10% FBS, with or without 100ug/ml of cycloheximide (or the same volume DMSO as control). Every 30 min after media exchange, aliquots of the media were collected and HiBiT activity was analysed with Nano-Glo HiBiT luciferase assay (Promega). This experiment was repeated 21 times.

### Purification of POLGARFin from cultured media and LC-MS analysis

To prepare the bait protein SNAP-LgBiT containing lysate, pcDNA3.4 SNAP-LgBiT plasmid was transfected into expi293F cells (Expifectamine 293 protocol). 48 h after transfection, 25×10^6^ cells expressing SNAP-LgBit were lysed in 5ml CCL buffer (Promega). The lysate aliquotes were stored at −80°C.

To prepare the conditioned media for POLGARFin purification, expi293F cells were transfected with pcDNA3.4 POLGARF-HiBiT (and pcDNA3.4 for negative control, Expifectamine 293 protocol). 48 h after transfection, the media was substituted for a fresh one, and transfected cells were grown for additional 48h. Then the cells were sedimented by centrifugation and the conditioned media was used for POLGARFin purification.

First, SNAP-LgBiT protein was loaded onto SNAP magnetic beads. For that, 40 μl of prewashed SNAP magnetic beads were incubated with 500 μl of SNAP-LgBiT lysate for 1.5 h at room temperature. Then the beads were washed in the CCL buffer supplemented with 0.5 M NaCl. Next, SNAP-LgBiT-modified magnetic beads were incubated with 10 mL of the conditioned medium for 30 min at room temperature with shaking. This procedure was repeated 5 times with fresh aliquots of the POLGARF-HiBiT-transfected cells medium. After the final incubation, the magnetic beads were washed in buffer A1000 (20 mM TrisHCl, 10% glycerol, 1mM DTT, EDTA, 1M KCl) and then washed 2 times in PBS (we have found that high salt wash do not disrupt association of HiBiT-containing proteins from SNAP-LgBiT). Bound proteins were eluted from the beads by incubation with 1xSDS-Loading dye for 5 min at 50°C and resolved on Tris-Tricine SDS PAGE. The gels were either stained with Coomassie (for mass spectrometry analysis) or transferred to nitrocellulose membrane and then analysed with HiBit-blot. Briefly, the membrane was incubated in TBS-T buffer for 30 min, followed by sequential incubation with the buffer, LgBiT protein, and Furimazine from Nano-Glo HiBiT luciferase assay (Promega) for 5 minutes followed by chemiluminescence detection on ChemiDoc XRS+ (BioRad).

For LC-MS sequencing, the HiBiT-containing protein was purified from the conditioned media using the SNAP-LgBiT bait protein, resolved on a Tris-Tricine gel, and stained with Coomassie. The protein band corresponding to the mobility of the POLGARFin-HiBiT protein in the blot experiment of the same eluate, was excised and analysed by LC-MS/MS. In-gel protein digestion was done as in (54) without protein reduction and Cys alkylation. Gel band was divided into six pieces, three were digested with GluC and the other three with GluC and trypsin. All samples were analysed by LC-MS in the same way as for the full-cell proteome analysis described above. Peptides were separated on an Acclaim Pepmap C18, 2 um column 75 um *150 mm using a 45 min gradient. Each sample was analyzed by a single LC-MS run.

All the obtained data files were analyzed together in PEAKS software against a Uniprot human database with a manually attached POLGARF sequence and the common contaminant database. Enzyme parameter was selected to be “Specified by each sample” with semispecific digestion. Samples digested with GluC were analyzed as digested with “GluC (bicarbonate)”. Samples digested with GluC and trypsin were analyzed with the digestion specificity combined from “GluC (bicarbonate)” and “Trypsin”. The results were filtered to PSM FDR 1%, which resulted in 2.1% peptide FDR and 0.2 % protein group FDR. After FDR filtering, 6 runs resulted in identification of 2617 PSM, 523 peptides and 53 protein groups, with most of the protein groups belonging to common contaminants.

### Riboseq data analysis

Publicly available Ribo-Seq/RNA-Seq datasets were downloaded from the NCBI Sequence Read Archive and converted to FASTQ format, see Table S3. After adapter and rRNA removal, reads below 25 nucleotides were discarded. Reads were then aligned to the GENCODE version 25 transcriptome using bowtie. Genomic positions of each read were determined and if aligning to only one genomic position were classed as unambiguous. A metagene profile was created for each Ribo-Seq read length from each dataset, centred around the annotated start codons. The distance between the start codon and the highest peak upstream of the start codon plus 3 nucleotides was used as an offset from the 5’ end of each read to infer the A-site. A triplet periodicity score was determined for each read length in each dataset by calculating the number of reads aligning to the same frame as the annotated CDS’s versus the number of reads in the −1/+1 frame. The triplet periodicity score was then defined as number of reads in the highest peak divided by the number of reads in the second highest peak, so that the read lengths with the strong periodicity would receive a score close to 1, while the read lengths with weak periodicity would receive a score close to 0.

### Data Presentation

The number of biological replicates for each experiment is indicated in each figure legend. For all Boxplots centre lines show the medians; box limits indicate the 25th and 75th percentiles as determined by R software; whiskers extend 1.5 times the interquartile range from the 25th and 75th percentiles, outliers are represented by dots. Genomic sequence alignments of 100 vertebrates were visually explored using CodAlignView (I. Jungreis, M. Lin and M. Kellis, manuscript in preparation) which was also used for generating Fig. 1C lower panels. The alignment shown in Fig. 3B was manually refined.

## Data Availability

The mass spectrometry proteomics data have been deposited to the ProteomeXchange Consortium via the PRIDE partner repository with the dataset identifier PXD016007

## Acknowledgments

INS, DEA and MSM acknowledge Russian Science Foundation (RSF16-14-10065); PVB is funded by SFI-HRB-Wellcome Trust Biomedical Research Partnership (Investigator Award in Science (210692/Z/18/Z); MALDI MS facility became available to us in the framework of the Moscow State University Development Program PNG 5.13; Funding for open access charge: Russian Science Foundation

