## Supplementary Information for "Unprecedentedly efficient CUG initiation of an overlapping reading frame in *POLG* mRNA yields novel protein POLGARF"

**Supplementary Table 1**. **POLGARF originated peptides detected with PATS.**

Eight tryptic peptide sequences of POLGARF were submitted to PATS (32) in order to identify particular datasets from HEK interactome study (33) where these peptides can be detected.

| POLGARF peptide sequence | Dataset ID | Bait |
|---|---|---|
| ---.AAAAAAAAAAAAAAATAASAAASAILGGR.--- | cs_b8855_CAMK2D.20847.20847.3 | CAMK2D |
| ---.AAAAAAAAAAAAAAATAASAAASAILGGR.--- | cs_j10057_CAMK2D.18684.18684.3 | CAMK2D |
| ---.AAAAAAAAAAAAAAATAASAAASAILGGR.--- | cs_b8855_CAMK2D.25107.25107.3 | CAMK2D |
| ---.AAAAAAAAAAAAAAATAASAAASAILGGR.--- | cs_q4994_CDK13.24081.24081.4 | CDK13 |
| ---.AAAAAAAAAAAAAAATAASAAASAILGGR.--- | cs_b6303_CDK3.19218.19218.4 | CDK3 |
| ---.AAAAAAAAAAAAAAATAASAAASAILGGR.--- | cs_j12279_CHCHD10.18793.18793.3 | CHCHD10 |
| ---.AAAAAAAAAAAAAAATAASAAASAILGGR.--- | cs_b13022_CLEC3A.24851.24851.3 | CLEC3A |
| ---.AAAAAAAAAAAAAAATAASAAASAILGGR.--- | cs_j9123_CRYAB.21338.21338.3 | CRYAB |
| ---.AAAAAAAAAAAAAAATAASAAASAILGGR.--- | cs_j6988_HNRNPA1.22831.22831.3 | HNRNPA1 |
| ---.AAAAAAAAAAAAAAATAASAAASAILGGR.--- | cs_b6142_HNRNPA1.21215.21215.3 | HNRNPA1 |

| | | |
|---|---|---|
| ---.AAAAAAAAAAAAAAATAASAAASAILGGR.--- | cs_b5359_KCTD17.20233.20233.3 | KCTD17 |
| ---.AAAAAAAAAAAAAAATAASAAASAILGGR.--- | cs_j6106_KCTD17.20598.20598.3 | KCTD17 |
| ---.AAAAAAAAAAAAAAATAASAAASAILGGR.--- | cs_q8280_KLHDC2.23818.23818.4 | KLHDC2 |
| ---.AAAAAAAAAAAAAAATAASAAASAILGGR.--- | cs_j12075_MAP2K2.21702.21702.2 | MAP2K2 |
| ---.AAAAAAAAAAAAAAATAASAAASAILGGR.--- | cs_j10965_NPM2.20787.20787.3 | NPM2 |
| ---.AAAAAAAAAAAAAAATAASAAASAILGGR.--- | cs_b5961_TIRAP.18987.18987.3 | TIRAP |
| ---.AAAAAAAAAAAAAAATAASAAASAILGGR.--- | cs_b5578_TMEM171.21238.21238.3 | TMEM171 |
| ---.AAAAAAAAAAAAAAATAASAAASAILGGR.--- | cs_j11048_TRIP13.19902.19902.3 | TRIP13 |
| ---.AAAAQPIGHPDALER.--- | cs_j10586_ALCAM.15890.15890.3 | ALCAM |
| ---.AAAAQPIGHPDALER.--- | cs_j13027_C1orf131.18647.18647.3 | C1orf131 |
| ---.AAAAQPIGHPDALER.--- | cs_j11428_C20orf114.10400.10400.3 | C20orf114 |
| ---.AAAAQPIGHPDALER.--- | cs_j10057_CAMK2D.5988.5988.3 | CAMK2D |
| ---.AAAAQPIGHPDALER.--- | cs_b8855_CAMK2D.6099.6099.3 | CAMK2D |
| ---.AAAAQPIGHPDALER.--- | cs_b8855_CAMK2D.6157.6157.2 | CAMK2D |
| ---.AAAAQPIGHPDALER.--- | cs_j9085_CDC20B.11650.11650.2 | CDC20B |
| ---.AAAAQPIGHPDALER.--- | cs_j12749_CDCA4.15371.15371.3 | CDCA4 |
| ---.AAAAQPIGHPDALER.--- | cs_b11228_CHCHD10.5486.5486.3 | CHCHD10 |
| ---.AAAAQPIGHPDALER.--- | cs_j12279_CHCHD10.5345.5345.3 | CHCHD10 |
| ---.AAAAQPIGHPDALER.--- | cs_b13022_CLEC3A.6877.6877.3 | CLEC3A |
| ---.AAAAQPIGHPDALER.--- | cs_b12828_CLEC3A.6001.6001.3 | CLEC3A |
| ---.AAAAQPIGHPDALER.--- | cs_q5682_CLEC4M.15058.15058.3 | CLEC4M |
| ---.AAAAQPIGHPDALER.--- | cs_j8140_DCAF4L1.12114.12114.3 | DCAF4L1 |
| ---.AAAAQPIGHPDALER.--- | cs_j8585_DDB2.12513.12513.3 | DDB2 |
| ---.AAAAQPIGHPDALER.--- | cs_j11233_DHX32.14164.14164.3 | DHX32 |
| ---.AAAAQPIGHPDALER.--- | cs_b10273_F9.13952.13952.3 | F9 |

| | | |
|---|---|---|
| ---.AAAAQPIGHPDALER.--- | cs_j5941_FHL3.5711.5711.3 | FHL3 |
| ---.AAAAQPIGHPDALER.--- | cs_b10379_GATAD2A.11144.11144.4 | GATAD2A |
| ---.AAAAQPIGHPDALER.--- | cs_j6880_HAGH.11335.11335.2 | HAGH |
| ---.AAAAQPIGHPDALER.--- | cs_b9320_HAVCR2.7294.7294.3 | HAVCR2 |
| ---.AAAAQPIGHPDALER.--- | cs_j10896_MGRN1.17168.17168.3 | MGRN1 |
| ---.AAAAQPIGHPDALER.--- | cs_b9791_NPM2.5928.5928.3 | NPM2 |
| ---.AAAAQPIGHPDALER.--- | cs_j10965_NPM2.6147.6147.3 | NPM2 |
| ---.AAAAQPIGHPDALER.--- | cs_b8377_P2RX1.14623.14623.3 | P2RX1 |
| ---.AAAAQPIGHPDALER.--- | cs_j7317_PEG10.5896.5896.3 | PEG10 |
| ---.AAAAQPIGHPDALER.--- | cs_b4647_PSME3.5472.5472.3 | PSME3 |
| ---.AAAAQPIGHPDALER.--- | cs_j11058_RNF135.15110.15110.3 | RNF135 |
| ---.AAAAQPIGHPDALER.--- | cs_q10121_SPACA3.22399.22399.3 | SPACA3 |
| ---.AAAAQPIGHPDALER.--- | cs_b12128_SSC5D.6081.6081.3 | SSC5D |
| ---.AAAAQPIGHPDALER.--- | cs_b12849_SSSCA1.5396.5396.3 | SSSCA1 |
| ---.AAAAQPIGHPDALER.--- | cs_q4961_TFG.4399.4399.3 | TFG |
| ---.AAAAQPIGHPDALER.--- | cs_q10216_TMEM213.5717.5717.3 | TMEM213 |
| ---.AAAAQPIGHPDALER.--- | cs_b10069_TNFRSF10A.6047.6047.3 | TNFRSF10A |
| ---.AAAAQPIGHPDALER.--- | cs_b11278_TRIM28.8946.8946.3 | TRIM28 |
| ---.AAAAQPIGHPDALER.--- | cs_j10886_TRIM29.14432.14432.3 | TRIM29 |
| ---.AAAAQPIGHPDALER.--- | cs_q8328_TRIP13.5873.5873.3 | TRIP13 |
| ---.AAAAQPIGHPDALER.--- | cs_j11048_TRIP13.5940.5940.3 | TRIP13 |
| ---.AAAAQPIGHPDALER.--- | cs_b5666_TSPAN5.6276.6276.3 | TSPAN5 |
| ---.AAAAQPIGHPDALER.--- | cs_b10186_WDR49.15088.15088.3 | WDR49 |
| ---.AGSSSGALGLQLRPR.--- | cs_b8932_BIN3.8961.8961.3 | BIN3 |
| ---.AGSSSGALGLQLRPR.--- | cs_j12830_CCDC59.10986.10986.2 | CCDC59 |

| | | |
|---|---|---|
| ---.AGSSSGALGLQLRPR.--- | cs_b12965_CDH8.11441.11441.4 | CDH8 |
| ---.AGSSSGALGLQLRPR.--- | cs_b6370_CHID1.12806.12806.2 | CHID1 |
| ---.AGSSSGALGLQLRPR.--- | cs_b12325_COBL.11688.11688.3 | COBL |
| ---.AGSSSGALGLQLRPR.--- | cs_j10834_CTPS2.5148.5148.3 | CTPS2 |
| ---.AGSSSGALGLQLRPR.--- | cs_j9237_EDC3.8135.8135.4 | EDC3 |
| ---.AGSSSGALGLQLRPR.--- | cs_b8092_EDC3.7109.7109.3 | EDC3 |
| ---.AGSSSGALGLQLRPR.--- | cs_b9320_HAVCR2.11223.11223.3 | HAVCR2 |
| ---.AGSSSGALGLQLRPR.--- | cs_b9320_HAVCR2.11200.11200.3 | HAVCR2 |
| ---.AGSSSGALGLQLRPR.--- | cs_q9743_HOXC10.9022.9022.2 | HOXC10 |
| ---.AGSSSGALGLQLRPR.--- | cs_b12842_IL28B.18406.18406.3 | IL28B |
| ---.AGSSSGALGLQLRPR.--- | cs_b8336_JMJD6.3034.3034.4 | JMJD6 |
| ---.AGSSSGALGLQLRPR.--- | cs_q5827_KIAA0415.17518.17518.3 | KIAA0415 |
| ---.AGSSSGALGLQLRPR.--- | cs_q8681_MOCS3.19021.19021.3 | MOCS3 |
| ---.AGSSSGALGLQLRPR.--- | cs_j12821_MRPS7.9784.9784.2 | MRPS7 |
| ---.AGSSSGALGLQLRPR.--- | cs_j13968_MUSK.9458.9458.2 | MUSK |
| ---.AGSSSGALGLQLRPR.--- | cs_b8713_OBP2A.12710.12710.2 | OBP2A |
| ---.AGSSSGALGLQLRPR.--- | cs_q5900_P2RX6.15858.15858.3 | P2RX6 |
| ---.AGSSSGALGLQLRPR.--- | cs_b9258_PNPLA2.18804.18804.3 | PNPLA2 |
| ---.AGSSSGALGLQLRPR.--- | cs_b7625_SUDS3.3524.3524.4 | SUDS3 |
| ---.AGSSSGALGLQLRPR.--- | cs_b10642_SYT16.11204.11204.3 | SYT16 |
| ---.AGSSSGALGLQLRPR.--- | cs_q5556_TADA1.8896.8896.3 | TADA1 |
| ---.AGSSSGALGLQLRPR.--- | cs_j5494_TP53RK.6812.6812.3 | TP53RK |
| ---.ALLLDQPAVAG.--- | cs_q4527_AKR1D1.11789.11789.2 | AKR1D1 |
| ---.ALLLDQPAVAG.--- | cs_j7359_DDX39A.10124.10124.2 | DDX39A |
| ---.ALLLDQPAVAG.--- | cs_b11016_FPR2.21858.21858.2 | FPR2 |

| | | |
|---|---|---|
| ---.ALLLDQPAVAG.--- | cs_j11797_LRRC33.15484.15484.2 | LRRC33 |
| ---.ALLLDQPAVAG.--- | cs_q6424_NAT8L.10424.10424.2 | NAT8L |
| ---.ALLLDQPAVAG.--- | cs_j12646_SLC25A43.9674.9674.2 | SLC25A43 |
| ---.GAAPAAPLR.--- | cs_b12201_C4orf6.8171.8171.2 | C4orf6 |
| ---.GAAPAAPLR.--- | cs_b12064_CDCP1.6161.6161.2 | CDCP1 |
| ---.GAAPAAPLR.--- | cs_j8621_CPB2.7356.7356.2 | CPB2 |
| ---.GAAPAAPLR.--- | cs_q4877_FBL.9064.9064.2 | FBL |
| ---.GAAPAAPLR.--- | cs_b4832_LDLRAP1.11276.11276.2 | LDLRAP1 |
| ---.GAAPAAPLR.--- | cs_j5406_LDLRAP1.11884.11884.2 | LDLRAP1 |
| ---.GAAPAAPLR.--- | cs_b9084_PDE1C.4715.4715.2 | PDE1C |
| ---.GAAPAAPLR.--- | cs_q10554_RANBP6.9691.9691.2 | RANBP6 |
| ---.GAAPAAPLR.--- | cs_q7821_RPE65.5284.5284.2 | RPE65 |
| ---.GAAPAAPLR.--- | cs_q10562_TLR5.8460.8460.2 | TLR5 |
| ---.GAAPAAPLR.--- | cs_q10562_TLR5.8489.8489.2 | TLR5 |
| ---.GAAPAAPLR.--- | cs_j11048_TRIP13.3915.3915.2 | TRIP13 |
| ---.GAAPAAPLR.--- | cs_q8039_WDTC1.5483.5483.2 | WDTC1 |
| ---.GAAPAAPLR.--- | cs_b6989_WIPI2.7125.7125.2 | WIPI2 |
| ---.GGPCTNHEPPALEEGGR.--- | cs_q7778_ACVRL1.14866.14866.2 | ACVRL1 |
| ---.GGPCTNHEPPALEEGGR.--- | cs_j9747_CAMKK1.17488.17488.2 | CAMKK1 |
| ---.GGPCTNHEPPALEEGGR.--- | cs_j13901_COMMD10.17499.17499.2 | COMMD10 |
| ---.GGPCTNHEPPALEEGGR.--- | cs_j6458_HLA-C.12386.12386.2 | HLA-C |
| ---.GGPCTNHEPPALEEGGR.--- | cs_j10300_MAU2.20438.20438.3 | MAU2 |
| ---.GGPCTNHEPPALEEGGR.--- | cs_q6713_NCOA5.11767.11767.2 | NCOA5 |
| ---.GGPCTNHEPPALEEGGR.--- | cs_j6174_SPRY1.11525.11525.2 | SPRY1 |
| ---.GGPCTNHEPPALEEGGR.--- | cs_q8206_STAM.8647.8647.2 | STAM |

| | | |
|---|---|---|
| ---.GGPCTNHEPPALEEGGR.--- | cs_j7197_TNFRSF11B.14598.14598.3 | TNFRSF11B |
| ---.GNLPHIGGGHIPLGLVFLVQPAAGGR.--- | cs_b8855_CAMK2D.20271.20271.4 | CAMK2D |
| ---.GNLPHIGGGHIPLGLVFLVQPAAGGR.--- | cs_j10057_CAMK2D.17076.17076.4 | CAMK2D |
| ---.GNLPHIGGGHIPLGLVFLVQPAAGGR.--- | cs_q8698_CYP2C9.16633.16633.4 | CYP2C9 |
| ---.GNLPHIGGGHIPLGLVFLVQPAAGGR.--- | cs_j7908_LTBR.19558.19558.4 | LTBR |
| ---.GNLPHIGGGHIPLGLVFLVQPAAGGR.--- | cs_j11048_TRIP13.17887.17887.4 | TRIP13 |
| --- .GQPGPALPPPGPEAEPALPGGGQLAVAGPAAPEAPGL GLGGGLDPVRPR.--- | cs_b9914_BHMT.21553.21553.7 | BHMT |
| --- .GQPGPALPPPGPEAEPALPGGGQLAVAGPAAPEAPGL GLGGGLDPVRPR.--- | cs_j9392_CNDP2.18838.18838.4 | CNDP2 |

**Supplementary Table 2**. **Primary and secondary antibodies used in this study**

| Company | Name | Catalogue N |
|---|---|---|
| Abcam, Cambridge, UK | Fibrillarin | ab4566 |
| EMD Millipore, MA | Mouse anti-Renilla | MAB4410 |
| Genscript, NJ | Custom made antibodies produced in rabbit against POLGARF peptide TNHEPPALEEGGRR | |
| LI-COR, NE | Donkey anti-mouse | 926-32212 |
| | Donkey anti-goat | 926-68024 |
| Promega, WI | Goat anti-Firefly | G745A |
| Santa-Cruz, Dallas, TX | RNA polymerase I (RPA194) | sc-48385 |
| Sigma, MO | $\alpha$–tubulin | T5168 |
| | Splicing Factor SC-35 | S4045 |
| | Mouse anti-rabbit IgG-peroxidase | A1949 |
| | Goat anti-mouse IgG-peroxidase | A0168 |
| | FLAG | F3165 |
| Thermo Fisher Scientific, MA | Goat anti-Rabbit IgG, Alexa Fluor 555 | A-21428 |
| | Donkey anti-Mouse IgG, Alexa Fluor 555 | A-31570 |
| | Goat anti-Mouse IgG, Alexa Fluor 647 | A-32728 |

**Supplementary table 3**. **Publicly available Riboseq datasets used in the analysis.**

| First Author | SRP Number | Year | PMID |
|---|---|---|---|
| Gawron | SRP065022 | 2016 | 26893308 |
| Gonzalez | SRP031501 | 2014 | 25122893 |
| Jakobsson | SRP102616 | 2017 | 28520920 |
| Battle | SRP047476 | 2015 | 25657249 |
| Xu | SRP052229 | 2016 | 26729373 |
| Park | SRP103009 | 2017 | 28494858 |
| Park | SRP072459 | 2016 | 27153541 |
| Werner | SRP048825 | 2015 | 26399832 |
| Andreev | SRP038695 | 2015 | 25621764 |
| Shi | SRP083699 | 2017 | 28106072 |
| Ji | SRP054971 | 2016 | 26900662, 26687005 |
| Elkon | SRP056200 | 2015 | 26538417 |
| Calviello | SRP063852 | 2016 | 26657557 |
| Guo | SRP031849 | 2014 | 25070500 |
| Kirchner | SRP065282 | 2017 | 28510592 |
| Iwasaki | SRP059825 | 2016 | 27309803 |
| Crappe | SRP042937 | 2015 | 25510491 |
| Sidrauski | SRP053402 | 2015 | 25719440 |
| Tirosh | SRP059531 | 2015 | 26599541 |
| Cenik | SRP055009 | 2015 | 26297486 |

| | | | |
|---|---|---|---|
| Zur | SRP064313 | 2016 | 26898226 |
| Zhang | SRP098797 | 2017 | 29170441 |
| Li | SRP114321 | 2017 | 29107537 |
| Oh | SRP060676 | 2016 | 27058758 |
| LoayzaPuch | SRP065530 | 2016 | 26878238 |
| Khajuria | SRP092068 | 2018 | 29551269 |

**Supplementary table 4**. **Publicly available proteomics datasets reanalysed for POLGARF search**

| published in | ID | source | description |
|---|---|---|---|
| Kulak et al., 2017 | PXD005141 | ProteomeXchange | Ultradeep proteomics with sample prefractionation for several cell lines. Jurkat cell line data was used. |
| Rieckmann et al., 2017 | PXD004352 | ProteomeXchange | Medium deep proteomics with sample prefractionation for immune cells. T4.naive, T4.EMRA, T8.naive and T8.EMRA data was used. |
| Bekker-Jensen et al., 2017 | PXD004452 | ProteomeXchange | Ultradeep proteomics with sample prefractionation for several cell lines. All present cell lines were reanalyzed. |
| Huttlin et al., 2017 | b8855_CAMK2D<br>b9320_HAVCR2<br>b9791_NPM2<br>b11228_CHCHD10<br>b12828_CLEC3A<br>j10057_CAMK2D<br>j10965_NPM2<br>j11048_TRIP13<br>j12279_CHCHD10<br>q7684_HAVCR2<br>q8328_TRIP13 | BioPlex | IP-MS experiments.<br>Baits from Supplementary Table 1 were taken for reanalysis if >=2 peptides were expected to be identified. |

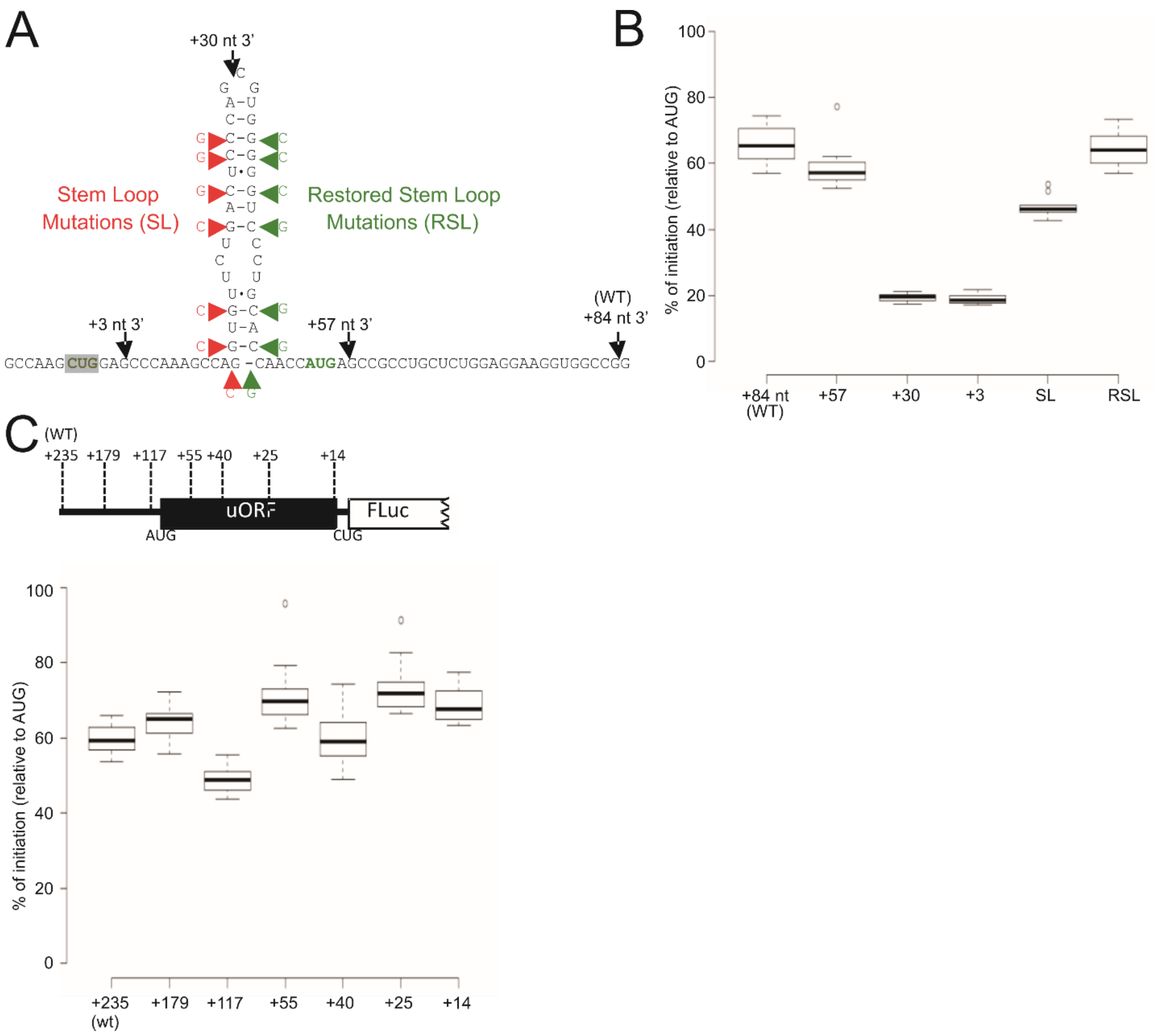


**Supplementary Figure 1**. **A.** Several previous studies reported that stable RNA secondary structures positioned approximately ~14 nt 3' of a poor context start codon can stimulate translation initiation (55). RNAFOLD (56) predicts a 35 nt RNA stem loop that starts 13 nt 3' of the *POLGARF* CUG initiation codon (notably, this stem loop is not conserved in many placental mammals). The positions of nested 3' deletions are indicated by black arrow heads, changes predicted to disrupt the stem-loop (SL) are indicated by red arrow heads and changes predicted to restore the stem-loop (RSL) are indicated by green arrow heads. The *POLGARF* initiation codon (CUG) is highlighted in grey and the *POLG* initiation codon (AUG) is in green font. **B.** Initiation efficiencies of firefly luciferase reporters fused to *POLGARF* 5' leaders with changes to 3' sequences as depicted in A. (n=12). **C.** Initiation efficiencies of firefly luciferase reporters fused to *POLGARF* 5' leaders with nested 5' deletions as indicated. (n=12).

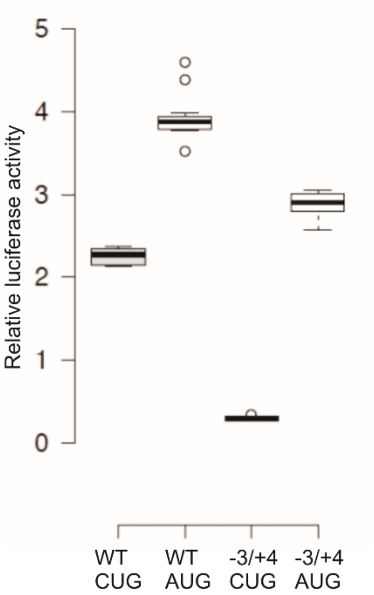


**Supplementary Figure 2**. Relative luciferase activities of firefly reporters fused to the *POLGARF* 5' leader. Shown are the wild-type CUG-initiated 5'leader (WT CUG), a CUG to AUG mutant (WT AUG), a -3 A-G plus +4 G-A mutant in the context of either CUG (CUG -3/+4) or AUG (AUG -3/+4)-initiated reporters. (n=12).

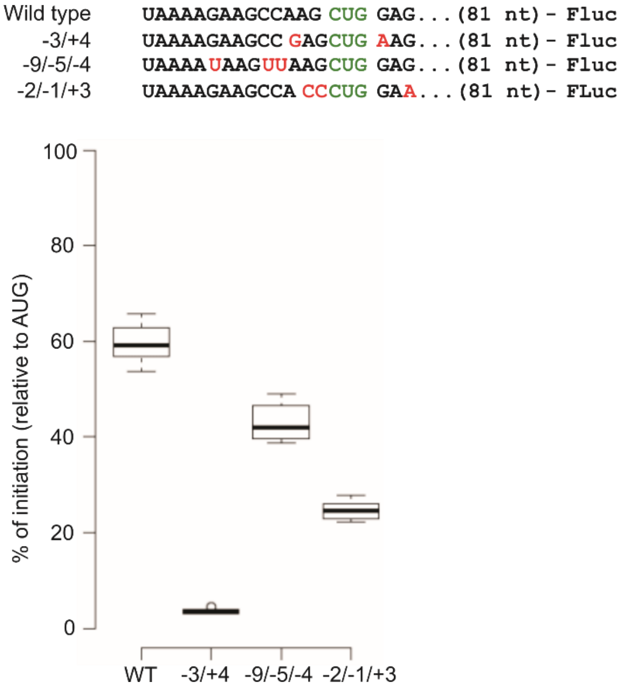


**Supplementary Figure 3**. Initiation efficiencies of firefly luciferase reporters fused to *POLGARF* 5’ leaders with changes to local context as indicated. (n=12).

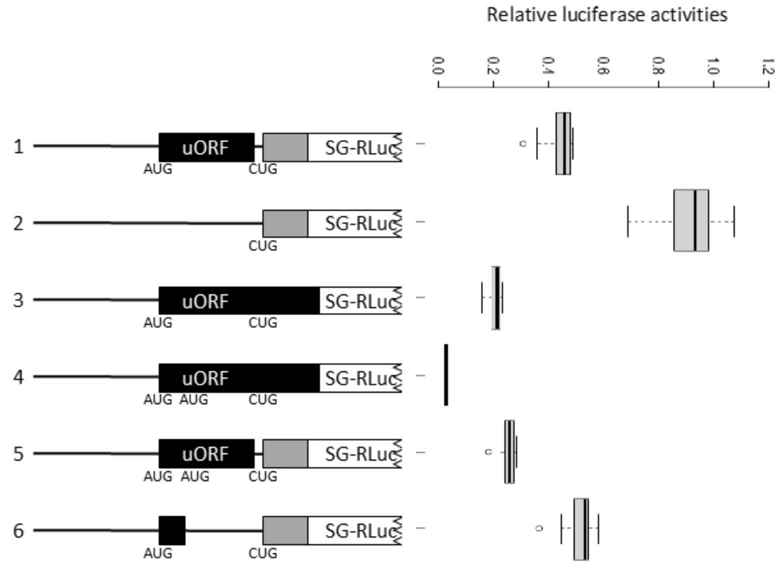


**Supplementary Figure 4**. Relative luciferase activities from -1 frame reporters fused to *POLG* 5' leaders with changes to the uORF. (n=12).

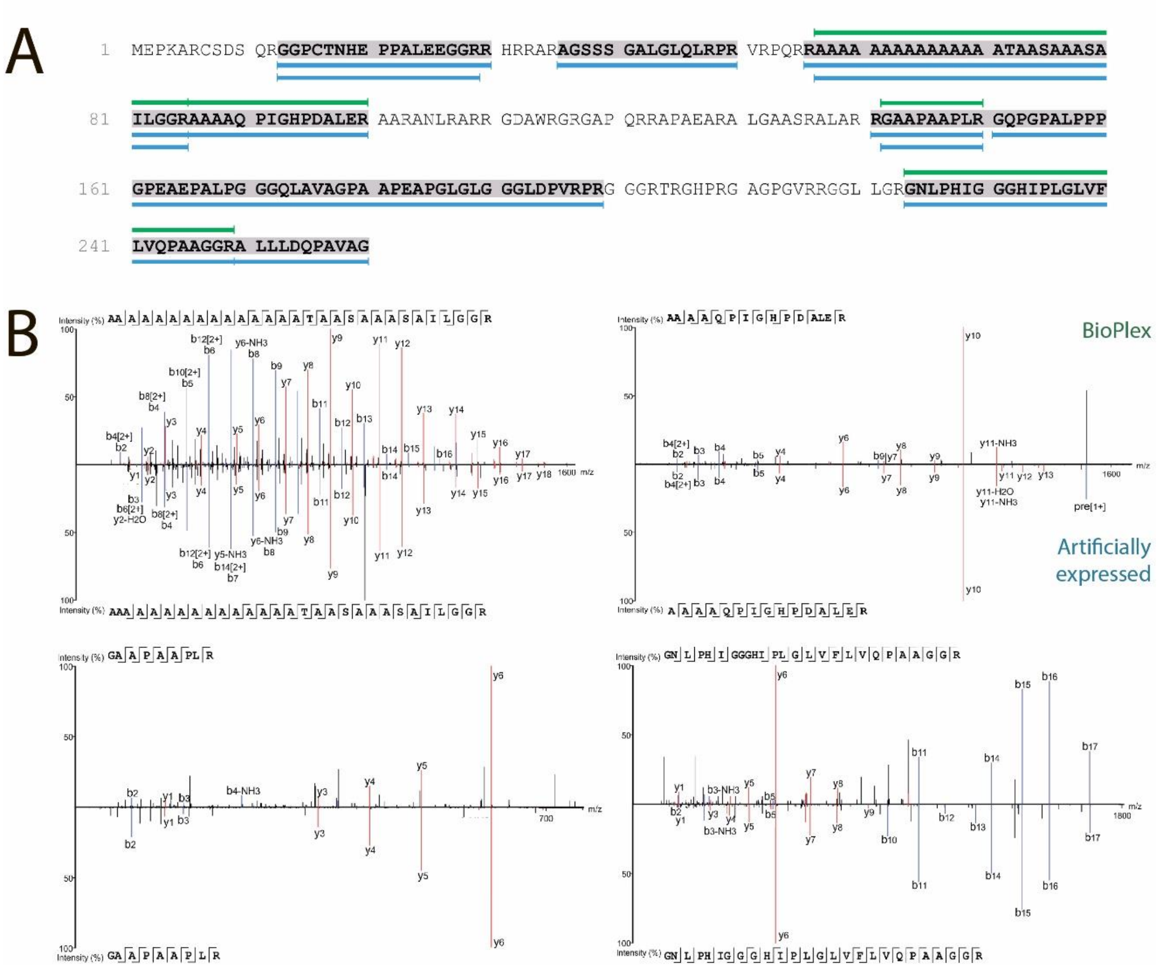


**Supplementary Figure 5.** Identification of POLGARF peptides in proteomic datasets. **A**. Coverage of peptides detected in Bioplex datasets (green) and peptides from lysates with overexpressed POLGARF (blue). **B**. Comparison of corresponding MS/MS spectra

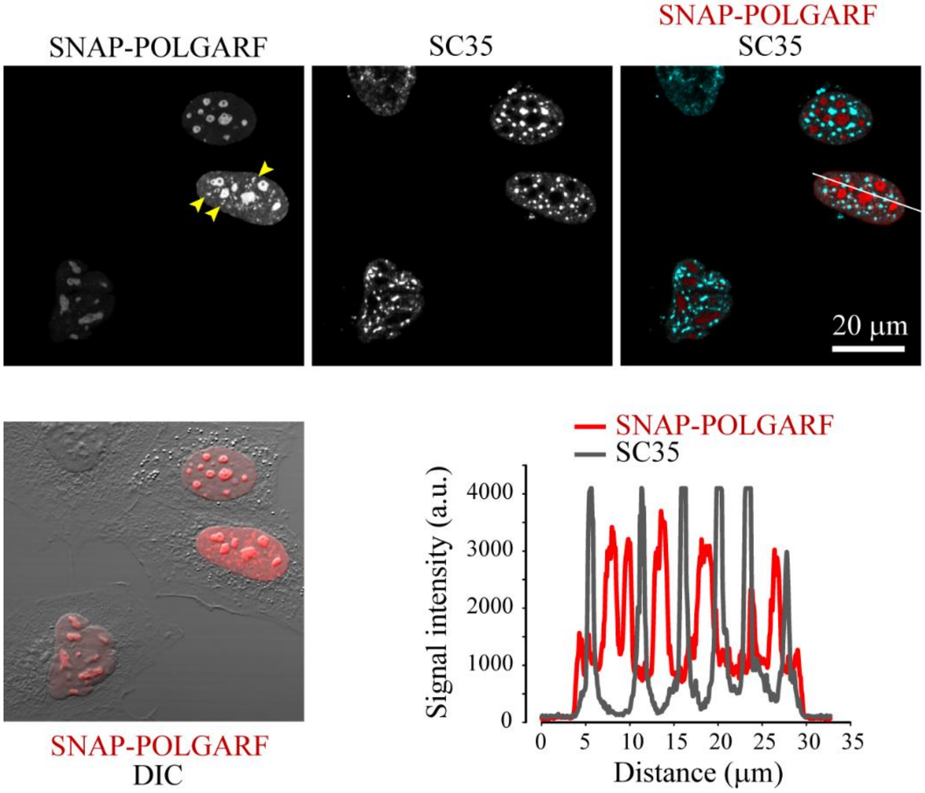


**Supplementary Figure 6.** Analysis of POLGARF and SC35 co-localization. Confocal fluorescence images of HEK293T cells transfected with pcDNA3.4 plasmid encoding SNAP-POLGARF represent stacks of 6 focal planes taken with a 0.5 μm step. SNAP was stained in live cells with SNAP-Cell® 647-SiR probe (excitable at 633 nm). Cells were fixed and stained with anti-SC35 antibodies conjugated with Alexa555 (excitable at 543 nm). Line profile analysis across a single cell (white line) demonstrates that non-nucleolar aggregates of POLGARF (marked by yellow arrowheads) do not co-localize with a pre-mRNA slicing factor SC35, suggesting that POLGARF is not involved in the early mRNA processing.

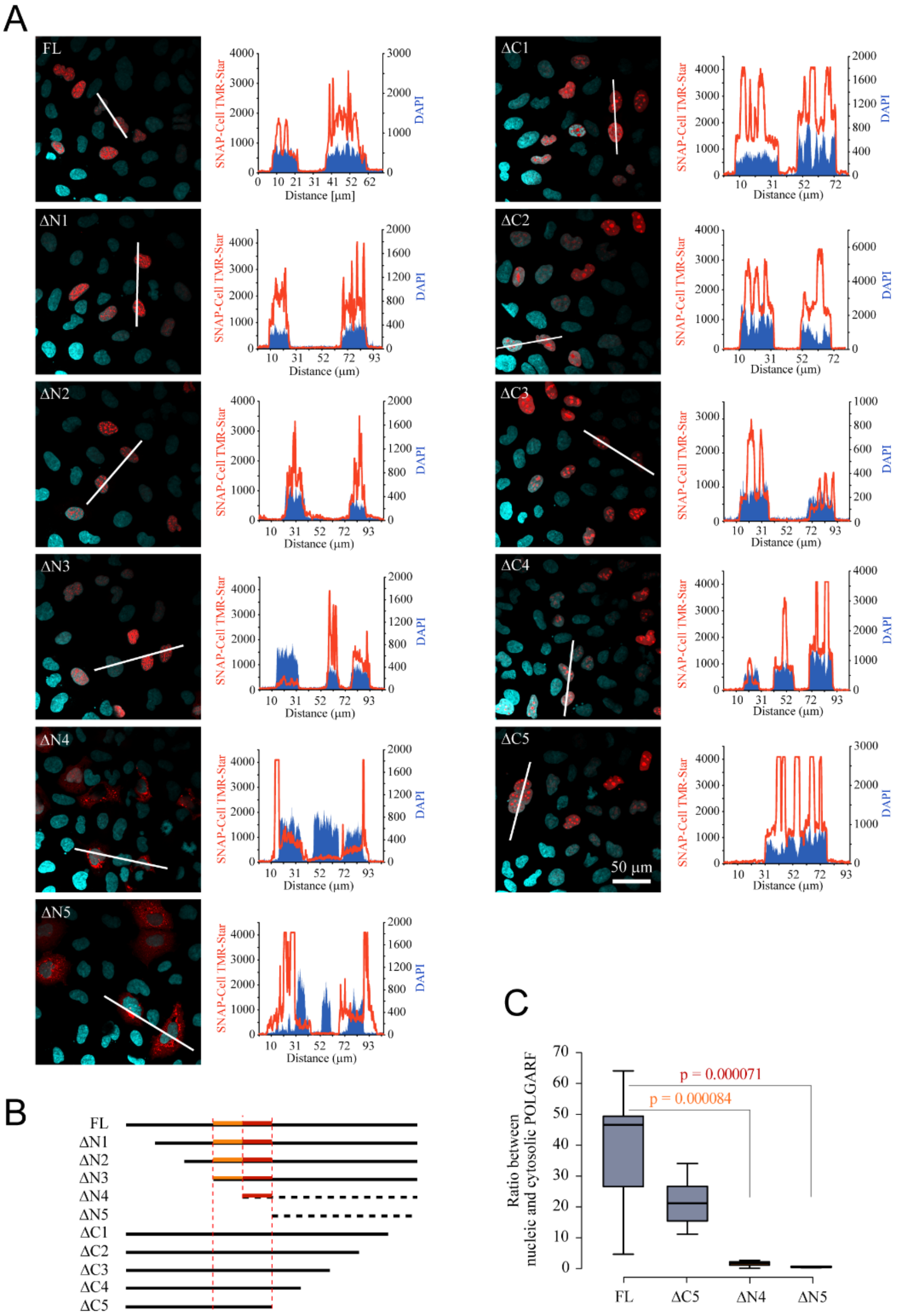


**Supplementary Figure 7**. Identification of the POLGARF fragment responsible for nuclear / nucleolar localization. **A**. Confocal images of HEK293T cells transfected with pcDNA3.4 plasmid encoding SNAP-POLGARF full length (FL) and its truncated versions shown in **B**. SNAP was stained in live cells with SNAP-Cell® TMR-Star probe (excitable at 543 nm). Cells were fixed and counterstained with DAPI. Images represent stacks of 8 focal planes taken with a 0.5 μm step. Line profile analysis across representative cells (white lines) is shown for each construct. **B**.

POLGARF fragments involved in nuclear / nucleolar localization are highlighted on the protein map in orange and red. **C.** Effects of truncation on the ratio between nuclear and cytosolic fractions of POLGARF; p-values (t-test) show dramatic effect of the two N-terminal fragments on the preference in POLGARF localization; $9 \leq n \leq 13$ cells.

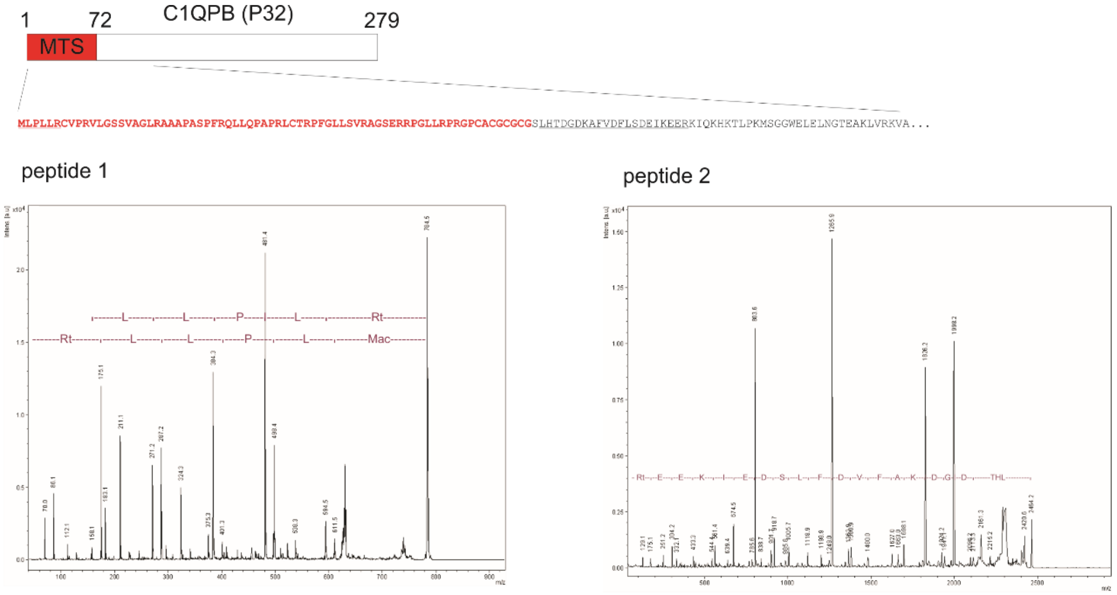


**Supplementary Figure 8**. MALDI TOF analysis of P32 from nucleolar fractions (extracted from lane 2 of the gel presented at Fig 4E) confirm the presence of an intact MTS sequence.

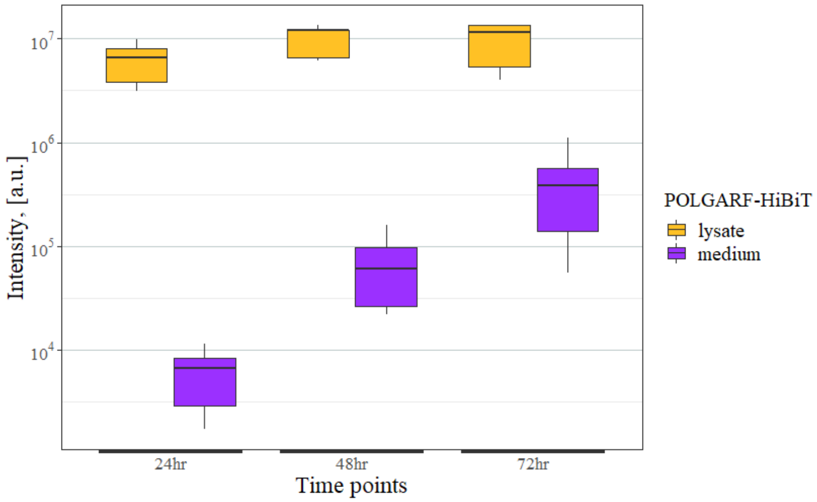


**Supplementary Figure 9**. Measurement of HiBiT activity in cell lysates and in cultured media 24, 48 and 72h after transfection of Hek293T cells with pcDNA3.4. POLGARF-HiBiT.

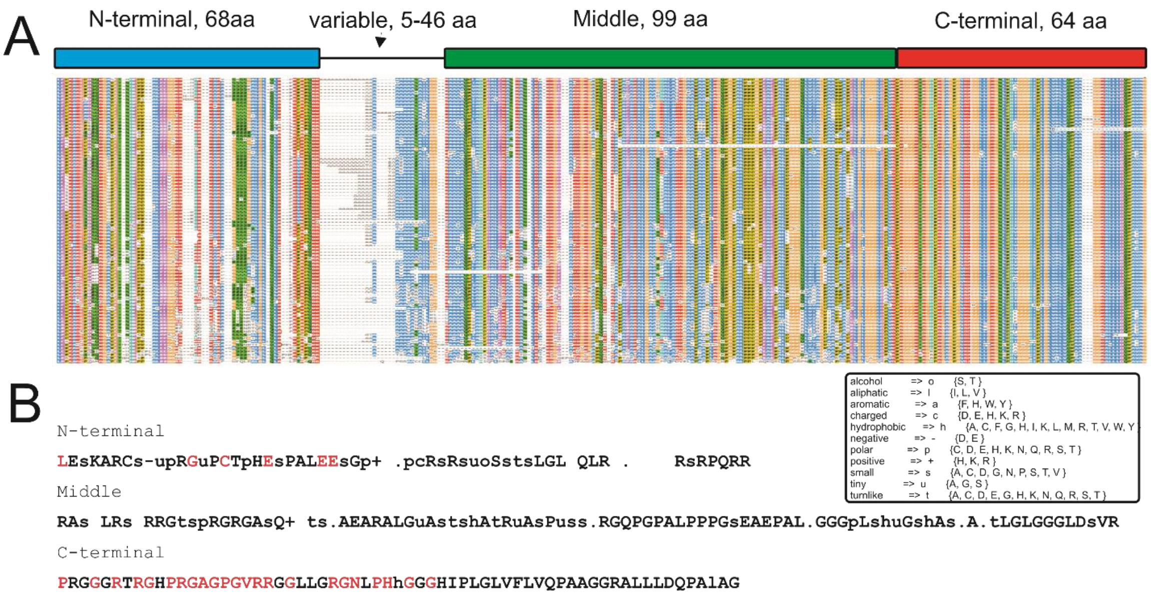


**Supplementary Figure 10.** Schematic organization of POLGARF protein. **A.** Schematic representation of POLGARF and multiple sequence alignment generated for 85 placental mammals. **B.** Consensus sequence of each part of POLGARF observed for 90% of sequences. 100% conserved amino acids are highlighted in red.
